# A brief breathing challenge reveals a cardiac correlate of generalized anxiety beyond resting physiology and interoceptive measures

**DOI:** 10.64898/2026.11.12.766286

**Authors:** Xindan Zhang, Huong Quynh Ngo, Keqing Shi, Timo L. Kvamme, Chengteng Ip, Juha Silvanto

## Abstract

Physiological markers measured at rest, such as heart rate variability, have proved relatively weak predictors of mental health symptoms. One potential reason is that resting measures are influenced by uncontrolled state factors, such as spontaneous breathing, whereas a standardized challenge holds these constant and may reveal regulatory differences not evident at rest. Here we asked whether respiratory sinus arrhythmia (RSA) during slow paced breathing, a challenge that strongly increases RSA, carries information about symptoms beyond resting RSA and interoceptive measures. In 207 young adults (mean age 23.6 years, SD = 3.1), electrocardiogram (ECG) was recorded at rest and during slow (6 breaths/min) and fast (20 breaths/min) paced breathing. Lower RSA during slow breathing, relative to that predicted by resting RSA, was associated with more severe generalized anxiety symptoms (β = −0.21, BF₁₀ = 13), also when depressive symptoms were controlled, whereas resting RSA was not. The association was concentrated among participants with at least moderate anxiety symptoms, whose resting RSA was typical but whose rest-adjusted slow-breathing RSA was 0.62 SD lower. Interoceptive accuracy, measured with heartbeat counting or heart rate discrimination, was unrelated to symptoms, self-report and slow-breathing RSA, and interoceptive questionnaires tracked distress shared across anxiety and depressive symptoms rather than generalized anxiety symptoms. At the group level, slow-breathing RSA thus carries information about generalized anxiety symptoms beyond resting RSA and self-reported interoception. Brief paced-breathing challenges may therefore increase the sensitivity of autonomic markers of anxiety.

## Introduction

Generalized anxiety is characterized by persistent worry and difficulty relaxing, and reduced activity of the vagus nerve, the main parasympathetic pathway that slows the heart, has long been proposed as one of its mechanisms (Thayer & Lane, 2000). Evidence from resting heart rate variability (HRV), the most common index of cardiac vagal activity, is mixed. HRV was lower than in controls in some case–control studies (Lyonfields et al., 1995; Pittig et al., 2013; Thayer et al., 1996) but not in others (Hammel et al., 2011; Kupats et al., 2018). Associations with symptom severity are weaker still (e.g. Óvári et al., 2026) and absent after control for depressive symptoms in a transdiagnostic clinical sample (Adolph et al., 2025). Part of the reason may lie in the measurement. Resting HRV depends on how fast and deeply a person happens to breathe (Grossman & Taylor, 2007), and people with generalized anxiety tend to breathe faster, which by itself lowers respiratory sinus arrhythmia without any change in vagal tone (Kollai & Kollai, 1992; but see Thayer et al., 1996). Resting measures also index only tonic vagal control, not how it responds when conditions change (Laborde et al., 2018). A more informative test may therefore be to assess how vagal control changes under challenge.

Challenge studies have mostly measured vagal withdrawal under stress, with weak and inconsistent results in generalized anxiety (Campbell & Wisco, 2021). Worry reduced vagal activity in individuals with and without the disorder alike (Hammel et al., 2011; Thayer et al., 1996), and blunted withdrawal has been linked mainly to depression (Adolph et al., 2025; Hamilton & Alloy, 2016; Kircanski et al., 2016). The opposite direction, how strongly vagal control responds when conditions favour rest, maps more directly onto difficulty relaxing but has received far less attention. Reduced autonomic flexibility was proposed early as a feature of chronic anxiety (Friedman & Thayer, 1998; Hoehn-Saric et al., 1989), and later models extended it to vagal control (Friedman, 2007; Thayer & Lane, 2000). These models, however, were tested chiefly through the prediction of low resting HRV. In the one study that examined range, patients tended to show a narrower range of vagally mediated HRV across baseline, relaxation and worry, but because their vagal activity was lower in every condition, a reduced range could not be separated from a low level (Thayer et al., 1996). Unpaced relaxation did not raise vagal activity above baseline (Thayer et al., 1996) and revealed only the group difference already present at rest (Pittig et al., 2013). Whether generalized anxiety involves a reduced capacity to increase vagal control above resting levels has therefore not been tested.

A stronger probe exists in clinical autonomic testing. Respiratory sinus arrhythmia (RSA) is the vagally mediated rise and fall of heart rate with breathing. Breathing at about six breaths per minute amplifies it, because this rate lies close to the resonance frequency of the cardiovascular system, at which breathing-related and baroreflex-driven fluctuations in heart rate reinforce each other (Laborde et al., 2022; Lehrer & Gevirtz, 2014). The resulting response is a well-validated test of cardiovagal function, abolished by vagal blockade and long used to detect autonomic neuropathy (Ewing et al., 1985; Shields, 2009). Yet it has rarely been applied to generalized anxiety. One small study (n = 55) found no difference between individuals with generalized anxiety symptoms and controls, but it scored the response only as normal or abnormal against clinical criteria and did not relate it to symptom severity (Kupats et al., 2018).

Here we tested whether generalized anxiety symptoms are related to RSA during resonance breathing, beyond resting RSA. We recorded the electrocardiogram in 207 young adults at rest and during slow (6 breaths/min) and fast (20 breaths/min) paced breathing, with fast breathing as an active control. RSA was quantified at the breathing frequency. Because responses to a challenge depend on their starting level (Berntson et al., 1994; Wilder, 1967), we indexed slow-breathing RSA relative to the value predicted by resting RSA. This measure, rest-adjusted slow-breathing RSA, reflects the level attained during resonance breathing rather than raw change from rest. Two accounts could link it to anxiety. If generalized anxiety involves a reduced capacity to raise vagal control above its resting level, the association should be specific to resonance breathing. If pacing instead simply indexes tonic vagal function more cleanly by removing variation in spontaneous breathing, fast breathing should show a similar association, although its smaller RSA makes this only a partial test. We predicted that rest-adjusted slow-breathing RSA, but not resting RSA or rest-adjusted fast-breathing RSA, would be associated with generalized anxiety symptoms, and we compared this association with that for depressive symptoms.

We also related this measure to interoception, the sensing of internal bodily signals, which has been proposed as a transdiagnostic mechanism in anxiety and depression (Khalsa et al., 2018). Self-reported interoception is consistently associated with symptoms (Desmedt et al., 2022) but corresponds poorly with interoceptive accuracy (Garfinkel et al., 2015) and resting cardiac physiology (Oliveira et al., 2025). It may therefore reflect beliefs about the body or general distress rather than bodily function (Banellis et al., 2026; Desmedt et al., 2022), and whether it tracks physiology under challenge is unknown. Interoceptive accuracy, in turn, was unrelated to mental health symptoms in the largest psychophysical study to date (Banellis et al., 2026), and its relation to the cardiovagal response to resonance breathing has not been tested.

To our knowledge, the present study is the first to relate RSA during resonance breathing to the severity of generalized anxiety symptoms. In addition, by measuring self-reported interoception, interoceptive accuracy and cardiac physiology in the same individuals, it asks whether these capture shared distress or variance specific to generalized anxiety.

## Methods

### Participants

The sample comprised 207 participants with a mean age of 23.6 years (SD = 3.1, range 18–35; 160 females). All participants provided written informed consent before testing and received financial compensation on completion. The study was approved by the University of Macau Research Ethics Committee (BSE-1794) and conducted in accordance with the Declaration of Helsinki. Exclusion criteria were self-reported cardiovascular, respiratory, neurological or psychiatric disorders and current use of medications. Participants were instructed to refrain from caffeine-containing beverages and alcohol on the day of testing and to obtain adequate sleep the night before. Sessions were conducted between 10:00 and 18:00. Recruitment aimed for approximately 200 participants, a target set before data collection began. A sensitivity analysis showed that the smallest analysis sample (n = 192) provided 80% power (α = .05, two-tailed) to detect a partial correlation of approximately .20 (f² = 0.04) for a single predictor added to a regression model with five predictors

### Design and Procedure

The study used a within-subjects design with three 5-min physiological conditions: rest, slow paced breathing at 6 breaths/min, and fast paced breathing at 20 breaths/min. Participants were seated throughout. The resting condition was always administered first to obtain a baseline before any breathing manipulation; participants sat looking at a fixation cross on the computer screen, remained physically relaxed, and breathed spontaneously. The order of the slow and fast conditions was counterbalanced across participants. Each condition was separated from the next by 2.5 min of unpaced rest. During paced breathing, participants followed a visual guide on a computer screen. A white rectangle moved up and down while the words inhale and exhale marked the corresponding respiratory phases. In the slow condition, each inhalation and exhalation lasted 5 s, giving a respiratory frequency of 0.10 Hz (6 breaths/min). This lies close to the resonance frequency of the cardiovascular system in most adults, although individual resonance frequencies vary (approximately 4.5–7 breaths/min; Lehrer & Gevirtz, 2014). In the fast condition, each phase lasted 1.5 s, giving a frequency of 0.33 Hz (20 breaths/min). The fast condition served as an active control with comparable pacing demands but outside the resonance range.

### Questionnaire Measures

Participants completed questionnaires assessing symptoms, alexithymia, childhood maltreatment and self-reported interoception, all administered in validated Chinese versions, and one of two cardiac interoceptive accuracy tasks. Questionnaires were completed in a separate block, either before or after the physiological protocol (counterbalanced across participants), with a 10-min break between the two parts. The interoceptive accuracy task was completed before the breathing protocol, and was followed by a 10 minute rest period.

#### Symptom measures

Generalized anxiety was assessed with the Generalized Anxiety Disorder-7 (GAD-7; Spitzer et al., 2006; Chinese version: He et al., 2010), a seven-item measure of generalized anxiety symptoms over the past 2 weeks. Scores range from 0 to 21, with scores of 10 or above indicating moderate or greater anxiety. Depressive symptoms were assessed with the Patient Health Questionnaire-9 (PHQ-9; range 0–27; Kroenke et al., 2001; Chinese version: Wang et al., 2014). State and trait anxiety were assessed with the six-item Chinese short forms of the State–Trait Anxiety Inventory (STAI-S-6 and STAI-T-6; range 6–24 each; Du et al., 2022; original: Spielberger et al., 1983).

#### Alexithymia

Alexithymia was assessed with the 24-item Perth Alexithymia Questionnaire (PAQ; Preece et al., 2018; Chinese version: Cai et al., 2024), rated on a seven-point scale (total range 24–168). The PAQ yields a total score and five subscales: negative and positive difficulty identifying feelings, negative and positive difficulty describing feelings, and general externally orientated thinking.

#### Self-reported interoception

Interoceptive sensibility was assessed with three questionnaires: 1) Multidimensional Assessment of Interoceptive Awareness, Version 2 (MAIA-2; Mehling et al., 2018; Chinese version MAIA-2C: Teng et al., 2022), which comprises eight subscales: Noticing, Not-Distracting, Not-Worrying, Attention Regulation, Emotional Awareness, Self-Regulation, Body Listening and Trusting. 2) The short form of the Body Perception Questionnaire (BPQ-SF; Cabrera et al., 2018; original: Porges, 1993; Chinese version: Wang et al., 2020), from which the body awareness and autonomic reactivity scales were scored. 3) the Interoceptive Accuracy Scale (IAS; Murphy et al., 2020; Chinese version: Lin et al., 2023), which assesses beliefs about one’s ability to perceive internal bodily signals accurately. Because the eight MAIA subscales do not form a single factor, a MAIA general score was computed as the mean of the six subscales that load on a common factor (Noticing, Attention Regulation, Emotional Awareness, Self-Regulation, Body Listening and Trusting; α = .82), with Not-Worrying and Not-Distracting analysed separately (Ferentzi et al., 2021). Self-report measures were not selected on their association with symptoms. Analyses used each questionnaire score, all 11 scores entered together, or the MAIA general score. Visual imagery vividness, measured with the Vividness of Visual Imagery Questionnaire (VVIQ; Marks, 1973), and the Childhood Trauma Questionnaire–Short Form (Bernstein et al., 2003; Chinese version: Zhao et al., 2005) was also administered but is not analysed here.

### Task-based Interoceptive accuracy measures

Participants completed either the heartbeat counting task (n = 107) or the Heart Rate Discrimination task (n = 91), both at rest with continuous ECG recording. The first 107 participants completed heartbeat counting and the subsequent 91 the Heart Rate Discrimination task. The remaining 9 participants did not complete an accuracy task because of technical problems. In the Heart Rate Discrimination task (Legrand et al., 2022), participants completed 60 interoceptive trials. Each trial began with a 5-s period in which participants attended to their heartbeat without touching their pulse. They then heard a sequence of five tones and judged whether the tones were faster or slower than their heart rate. The tone rate differed from the participant’s mean heart rate during the attention period by an amount set adaptively by a Bayesian psychophysical procedure (Psi). Of the 60 trials, 48 were adaptive Psi trials and 12 (20%) were catch trials with large differences between tone rate and heart rate. After each judgement, participants rated their confidence. Only the interoceptive condition was administered. Interoceptive accuracy was indexed by the absolute value of the perceptual threshold, that is, its distance from zero, with smaller values indicating greater accuracy, and precision by the slope of the psychometric function.

In the heartbeat counting task (Schandry, 1981), participants silently counted their heartbeats during six intervals (25, 30, 35, 40, 45 and 50 s), presented in random order, without taking their pulse. They were instructed to report only heartbeats they actually felt and not to estimate. The number of R peaks in each interval was taken from the ECG. Accuracy was computed as the mean across intervals of 1 − |recorded − counted| / recorded (Garfinkel et al., 2015). Heartbeat counting has been criticized because performance can reflect estimation based on elapsed time or knowledge of one’s heart rate rather than perception of heartbeats (Desmedt et al., 2020; Zamariola et al., 2018). We included it because it remains the most widely used measure of cardiac interoceptive accuracy, allowing comparison with previous work, and used instructions that discourage estimation, which improve its validity (Desmedt et al., 2020). The Heart Rate Discrimination task, used in the later testing period, separates perceptual accuracy from response bias and is less susceptible to these problems (Legrand et al., 2022). Because heartbeat counting and discrimination-based measures correlate only weakly (Hickman et al., 2020), each task was analysed separately. A pooled score, standardized within each task, was also analysed.

### Physiological Data Acquisition and Preprocessing

Electrocardiogram (ECG) and respiration were recorded continuously at 2000 Hz with a BIOPAC MP160 system equipped with a BioNomadix RSPEC-R module and AcqKnowledge software (BIOPAC Systems, Goleta, CA). Three ECG electrodes were placed on the left and right sides of the rib cage and below the right clavicle. Respiration was measured with a respiratory belt positioned around the thorax. The belt signal was not calibrated to volume, so respiratory amplitude indexes relative breathing depth rather than tidal volume.

Raw ECG signals were high-pass filtered (zero-phase second-order Butterworth, 0.5 Hz cutoff) to remove baseline drift, followed by five-sample smoothing. QRS complexes were enhanced by discrete wavelet decomposition (Symlet-4, six levels), reconstructing the signal from weighted absolute detail coefficients at levels 4, 5 and 6 (weights 0.3, 1.0 and 1.5). The reconstructed signal was low-pass filtered at 40 Hz and normalized. R peaks were detected with an adaptive algorithm (minimum inter-peak distance 0.40 × sampling frequency; minimum prominence 30% of the difference between the 80th and 97th percentiles of the enhanced signal amplitude). Candidate peaks were refined by identifying local ECG maxima within ±40 ms and were screened with an adaptive amplitude threshold set at 60% of the local 85th-percentile R-peak height within a 30-s window. For three participants, these settings were adjusted after visual inspection of the ECG and applied to all three conditions. The amplitude screening was omitted for all three, and for two the minimum inter-peak distance was increased to 0.65 × sampling frequency to prevent T waves being detected as R peaks. Artefactual R–R intervals were identified using interval-duration, successive-difference, median-absolute-deviation and R-peak-amplitude criteria (see Data Quality and Exclusions), and flagged intervals and their immediate neighbours were excluded. The remaining R–R intervals were interpolated at 4 Hz using piecewise cubic Hermite interpolation, and heart-period power was estimated with Welch’s method (64-s Hamming windows, 50% overlap).

Respiratory signals were linearly detrended and low-pass filtered at 1 Hz. They were used to verify breathing compliance, estimate respiratory frequency, and quantify respiratory amplitude as the peak-to-trough excursion of each respiratory cycle. Respiratory frequency was estimated from the dominant spectral peak of the respiratory signal within condition-specific ranges (0.07–0.13 Hz for slow breathing and 0.23–0.43 Hz for fast breathing). Respiratory amplitude was summarized as the median peak-to-trough excursion across detected cycles. For each condition, a 5-min analysis window was used. Breathing rate was computed from the detected respiratory cycles.

### Physiological Measures

#### Respiratory sinus arrhythmia

The primary index was respiratory sinus arrhythmia (RSA), quantified as the natural logarithm of heart-period power in a band centred on the breathing frequency of each condition, spanning approximately ±30% of that frequency. The bands were 0.07–0.13 Hz for slow breathing and approximately 0.23–0.43 Hz for fast breathing, centred on the instructed frequencies of 0.10 and 0.33 Hz; for the resting condition, the band was centred on each participant’s spontaneous respiratory frequency. Respiration-centred bands were used rather than the conventional high-frequency band (0.15–0.40 Hz) as breathing at 6 breaths/min falls below that band, and RSA at the paced frequency isolates the resonance oscillation caused by slow breathing.

#### Rest-adjusted slow-breathing *RSA*

Rest-adjusted slow-breathing RSA. Slow-breathing RSA was indexed relative to resting RSA as the residual from a linear regression of slow-breathing RSA on resting RSA, that is, the deviation of each participant’s slow-breathing RSA from the value predicted by their resting RSA. This index is equivalent to raw slow-breathing RSA with resting RSA as a covariate; in models that also include resting RSA its coefficient is identical to that of raw slow-breathing RSA, and by definition it is uncorrelated with resting RSA. Residualized scores were used instead of simple differences because difference scores are correlated with baseline and susceptible to regression to the mean (Berntson et al., 1994). A rest-adjusted fast-breathing index was used as a control.

#### Heart rate

Following current guidelines (Quigley et al., 2024), mean heart rate was analysed alongside RSA to distinguish changes in cardiac level from changes in the respiratory oscillation. Mean heart rate was calculated as 60 divided by the mean of the retained R–R intervals (in seconds). Heart-rate slowing was defined as the residual of slow-breathing heart rate after linear regression on resting heart rate, sign-reversed so that higher values indicate greater slowing. Fast-breathing heart-rate change was computed analogously.

#### Other heart rate variability indices

The root mean square of successive differences between normal R–R intervals (RMSSD) and the standard deviation of normal R–R intervals (SDNN) were computed for each condition and log-transformed. For RMSSD, successive differences were calculated only between originally consecutive R–R intervals that both survived artefact screening. Rest-adjusted slow-breathing indices based on RMSSD and SDNN, computed in the same way as for RSA, were used in robustness analyses.

### Data Quality and Exclusions

Exclusion criteria were applied to the physiological data before any association with questionnaire measures was examined. Data quality was assessed in two stages. First, recordings were screened for artefacts before any statistical outlier treatment. R–R intervals were flagged as artefactual if their duration fell outside 0.30–2.00 s, if the absolute difference from the preceding interval exceeded 500 ms or 50% of the preceding interval, or if they deviated by more than five scaled median absolute deviations from the median R–R interval. R peaks whose amplitude deviated by more than 45% from the mean R-peak amplitude within the condition were also flagged, together with their R–R intervals. Flagged intervals and their immediate neighbours were excluded from heart rate variability calculations.

For each condition, an artefact ratio was calculated as the proportion of originally detected R– R intervals removed by this procedure. A condition was excluded if this ratio exceeded 0.30, if fewer than 100 beats were detected, or if heart-period power exceeded 10⁵ ms². That power threshold is an order of magnitude above normative values for short-term recordings in healthy adults (Nunan et al., 2010) and above the largest value observed in this sample during slow breathing (≈ 1.5 × 10⁴ ms²), so values beyond it indicate artefact. One participant with frequent premature ventricular contractions was excluded from all heart rate variability analyses. One participant had no usable resting recording, and artefact screening removed a further four recordings (two slow-breathing and two fast-breathing). Second, RSA values more than 3 SD from the condition mean were excluded iteratively within each condition (two resting, two slow-breathing and three fast-breathing values). Because the choice of outlier treatment, HRV metric and covariate set could influence the results, we report a specification curve across these choices (Supplementary Fig. 1), and sensitivity analyses with alternative outlier treatments are reported in the Supplementary Materials. Because of these exclusions and missing questionnaire data, sample sizes for the main analyses ranged from 192 to 199. Data quality did not differ by anxiety level: artefact ratios were unrelated to GAD-7 scores in any condition (Spearman ρs = −.01 to .06, ps ≥ .40), and the proportion of excluded recordings did not differ between participants below and at or above the threshold for moderate generalized anxiety (GAD-7 ≥ 10; Fisher’s exact ps ≥ .26).

### Statistical Analysis

The primary analysis tested whether rest-adjusted slow-breathing RSA predicted generalized anxiety symptoms beyond resting RSA, with rest-adjusted fast-breathing RSA as a control. These were the threshold-group comparisons, condition × anxiety interactions, the decomposition of symptom variance, and the heart-rate and phase-coupling analyses.

Analyses were conducted in Python 3 (statsmodels, scikit-learn). Continuous variables were standardized, so coefficients are standardized (β). Unless stated otherwise, models were adjusted for age, gender and resting breathing rate, which influences resting RSA. Tests were two-tailed (α = .05). For key tests, default Bayes factors compared regression models with and without the predictor of interest (Jeffreys–Zellner–Siow prior, r = 0.354; Rouder & Morey, 2012). BF₁₀ values above 3 and 10 were interpreted as moderate and strong evidence for an effect, BF₀₁ values above 3 and 10 as moderate and strong evidence for its absence, and values between 1/3 and 3 as inconclusive. Robustness to prior width was assessed at r = 0.5 and 0.707 (Supplementary Table 5).

#### Main analyses

A repeated-measures ANOVA with Bonferroni-corrected paired t tests verified that the breathing manipulation altered RSA. Each symptom score (GAD-7, PHQ-9, STAI-state, STAI-trait and PAQ) was regressed on resting RSA and rest-adjusted slow-breathing RSA. The variance added by the rest-adjusted index (ΔR²) was tested with an F test, and the model was repeated with rest-adjusted fast-breathing RSA. Specificity to generalized anxiety was tested in two ways. First, the GAD-7 and PHQ-9 coefficients were compared in participants with both measures (bootstrap CI, 5,000 resamples). Second, each measure was entered as a covariate in the model for the other.

#### Robustness

For GAD-7 scores, respiratory amplitude and mean heart period (rest and slow breathing) were added as covariates. Alternative outlier treatments were applied: no trimming, winsorizing at the 1st and 99th percentiles, rank-based regression, and exclusion beyond 3.5 median absolute deviations. Influence was assessed with Cook’s distance. A specification curve crossed six outlier treatments (the main 3-SD trimming, the four alternatives and iterative 2.5-SD trimming) with four covariate sets (none; age and gender; plus resting breathing rate; plus respiratory amplitude). This gave 24 specifications per outcome (Supplementary Fig. 1), and the curve was repeated with RMSSD and SDNN in place of RSA. Raw slow-breathing RSA and simple difference scores (slow minus resting RSA) were also tested.

#### Self-report and physiology

Each of the 11 interoception scales (IAS, two BPQ scales and eight MAIA subscales) was regressed on resting RSA and rest-adjusted slow-breathing RSA, with Benjamini–Hochberg false discovery rate (FDR) correction across the 22 tests. To test whether the GAD-7 association was independent of self-report, GAD-7 scores were regressed on both RSA indices together with each scale in turn and with all 11 together. The unique and shared contributions of physiology (both RSA indices) and self-report (all 11 scales) to GAD-7 and PHQ-9 scores were estimated with hierarchical regressions entering the two blocks in both orders. These are reported as adjusted R² because the self-report block contained 11 predictors. Interactions between rest-adjusted slow-breathing RSA and self-report were tested for the MAIA general score and for each scale separately (FDR-corrected).

#### Decomposition of symptom variance

A one-factor model of GAD-7, PHQ-9, STAI-state and STAI-trait scores (maximum likelihood; regression-method factor scores) defined a general distress factor. GAD-7 and PHQ-9 residuals after regression on this factor defined anxiety- and depression-specific components. Each component was regressed simultaneously on resting RSA, rest-adjusted slow-breathing RSA and the MAIA general score, and the analysis was then repeated with each questionnaire score in place of the MAIA general score.

#### Interoceptive accuracy

Performance was coded so that higher values indicate greater accuracy (heartbeat counting accuracy; negative absolute HRD threshold). Each task was analysed separately, and scores standardized within task were also pooled, with task as a covariate. Accuracy was tested against symptom scores, the symptom components, resting RSA, rest-adjusted slow- and fast-breathing RSA and each self-report scale.

These models were adjusted for age and gender, plus resting breathing rate for physiological outcomes. Because heartbeat counting performance depends on heart rate, the association between counting accuracy and resting RSA was also tested with resting heart rate controlled. FDR correction was applied across the 11 self-report scales and, for the HRD task, across the 16 tests of accuracy and precision against the symptom and physiological outcomes. Because not all participants completed an accuracy task, these models were run separately from the main analyses. HRD precision (slope) was analysed as a secondary index, and HRD analyses were repeated excluding participants with lapse rates above .20.

#### Heart-rate analyses

Heart-rate slowing was tested against each interoception scale (FDR-corrected) and each symptom measure. Associations surviving correction were tested for robustness to respiratory amplitude, RSA and anxiety. The main GAD-7 model was also repeated with heart-rate slowing as a covariate.

#### Other exploratory analyses

Rest-adjusted slow-breathing RSA was compared between participants scoring at or above versus below the threshold for moderate symptoms (≥ 10) on the GAD-7 and PHQ-9, adjusting for resting RSA and covariates, and across alternative GAD-7 thresholds from 8 to 12. Condition order was tested as a predictor of rest-adjusted slow-breathing RSA and as a moderator of the slow- and fast-breathing associations with GAD-7 scores. Whether the anxiety association differed between conditions was tested with linear mixed models (random intercept per participant). These included condition, anxiety (continuous GAD-7 score or threshold group), their interaction and covariates. They compared rest with slow breathing and fast with slow breathing, with RSA in raw units and standardized within condition. Participants at or above versus below the GAD-7 threshold were compared on respiratory amplitude and rate in each condition, the change in respiratory amplitude from rest to slow breathing (log ratio) and heart-rate change from rest.

The amplitude change was also added to the main model. RSA in each condition was summarized by GAD-7 band (0–4, 5–9, 10–14 and 15–21). The main model was repeated with STAI-trait scores as a covariate, in participants below the GAD-7 threshold only, and excluding participants whose resting breathing rate was below 0.15 Hz (9 breaths/min), where the resting band overlaps the low-frequency range. Finally, cardiorespiratory phase coupling was quantified in each condition as the phase-locking value, PLV = |mean(exp(i(φ_resp − φ_hp)))|. The instantaneous phases came from Hilbert transforms of the heart-period and respiratory signals, band-pass filtered within the same respiration-centred band as RSA. PLV ranges from 0 (no synchronization) to 1 (perfect synchronization). Fisher-transformed PLV values were regressed on GAD-7 scores, adjusting for resting coupling and covariates, and with PHQ-9 scores, respiratory amplitude or rest-adjusted slow-breathing RSA as additional covariates.

## Results

Descriptive statistics for all questionnaire, interoceptive accuracy and physiological measures are given in Table 1, and correlations among the main variables in Supplementary Table 1. Symptom levels were low to moderate on average (GAD-7: M = 5.99, SD = 4.29; PHQ-9: M = 5.91, SD = 3.75). In the full sample, 34 participants (16.5%) scored in the moderate range or above (≥ 10) on the GAD-7 and 32 (15.6%) on the PHQ-9; numbers in each analysis sample are given below.

**Table 1.** Descriptive Statistics for Questionnaire, Interoceptive Accuracy and Physiological Measures.

| Measure | n | M (SD) |  |
|---|---|---|---|
| *A. Questionnaires* |  |  |  |
| GAD-7 | 206 | 5.99 (4.29) |  |
| PHQ-9 | 205 | 5.91 (3.75) |  |
| STAI-state (short form) | 199 | 11.29 (3.48) |  |
| STAI-trait (short form) | 199 | 11.92 (3.40) |  |
| PAQ total | 204 | 63.17 (23.40) |  |
| Negative difficulty identifying feelings | 204 | 11.20 (5.03) |  |
| Positive difficulty identifying feelings | 204 | 9.44 (4.29) |  |
| Negative difficulty describing feelings | 204 | 12.23 (5.34) |  |
| Positive difficulty describing feelings | 204 | 11.06 (5.12) |  |
| General externally orientated thinking | 204 | 19.24 (8.81) |  |
| IAS | 205 | 77.73 (9.07) |  |
| BPQ body awareness | 205 | 57.20 (12.12) |  |
| BPQ autonomic reactivity | 205 | 37.89 (10.26) |  |
| MAIA Noticing | 205 | 11.83 (3.40) |  |
| MAIA Not-Distracting | 205 | 15.10 (5.95) |  |
| MAIA Not-Worrying | 205 | 11.22 (4.61) |  |
| MAIA Attention Regulation | 205 | 19.61 (6.12) |  |
| MAIA Emotional Awareness | 205 | 16.77 (4.34) |  |
| MAIA Self-Regulation | 205 | 11.33 (3.74) |  |
| MAIA Body Listening | 205 | 7.93 (3.36) |  |
| MAIA Trusting | 205 | 9.94 (2.64) |  |
| *B. Interoceptive accuracy* |  |  |  |
| Heartbeat counting accuracy (0–1) | 107 | 0.68 (0.15) |  |
| HRD threshold (bias, bpm) | 91 | −6.95 (14.85) |  |
| HRD absolute threshold (bpm) | 91 | 13.27 (9.58) |  |
| HRD slope | 91 | 8.55 (2.65) |  |
| *C. Physiological measures by condition* |  |  |  |
| Measure | Rest | Slow breathing | Fast breathing |
|  | *M (SD)* | *M (SD)* | *M (SD)* |
| RSA (ln ms²) | 5.66 (1.13) | 8.23 (0.64) | 5.38 (1.01) |
| Log-RMSSD (ln ms) | 3.34 (0.48) | 3.70 (0.37) | 3.36 (0.54) |
| Log-SDNN (ln ms) | 3.83 (0.38) | 4.37 (0.28) | 3.82 (0.36) |
| Heart rate (beats/min) | 78.7 (10.0) | 76.7 (8.9) | 80.2 (10.9) |
| Breathing rate (breaths/min) | 14.89 (3.41) | 6.00 (0.07) | 19.94 (0.23) |
| Respiratory amplitude (a.u.) | 2.60 (1.97) | 6.89 (3.95) | 3.83 (2.71) |
| Phase coupling (PLV) | 0.83 (0.16) | 0.95 (0.06) | 0.90 (0.10) |
| n (RSA / other cardiac / respiration) | 203 / 205 / 206 | 202 / 204 / 207 | 201 / 204 / 207 |

*Note.* PAQ = Perth Alexithymia Questionnaire; IAS = Interoceptive Accuracy Scale; BPQ = Body Perception Questionnaire (autonomic reactivity = supradiaphragmatic + subdiaphragmatic); MAIA = Multidimensional Assessment of Interoceptive Awareness, Version 2 (subscale sums); HRD = Heart Rate Discrimination. Heartbeat counting and HRD were completed by different participants.
*Note.* Cardiac measures are after artefact screening; RSA values are additionally after iterative 3-SD trimming, as in the main analyses. RSA = natural log of heart-period power in a band centred on each condition's breathing frequency; RMSSD and SDNN were log-transformed. Breathing rate was estimated from respiratory cycles. Respiratory amplitude is in arbitrary units (uncalibrated belt). PLV = phase-locking value between heart period and respiration.

### Slow breathing inceased respiratory sinus arrhythmia

Participants adhered closely to the protocol (breathing rate 6.00 breaths/min, SD = 0.07, during slow breathing and 19.94, SD = 0.23, during fast breathing). RSA differed across conditions (F(2, 392) = 1091, p < .001, n = 197). Slow breathing increased RSA relative to rest by 2.55 ln units (*d*z = 2.55) and relative to fast breathing by 2.85 ln units (*d*z = 3.40; both Bonferroni-corrected ps < .001; Fig. 1a); fast-breathing RSA was slightly lower than resting RSA (0.30 ln units, *d*z = 0.31, p < .001).

**Figure 1.**
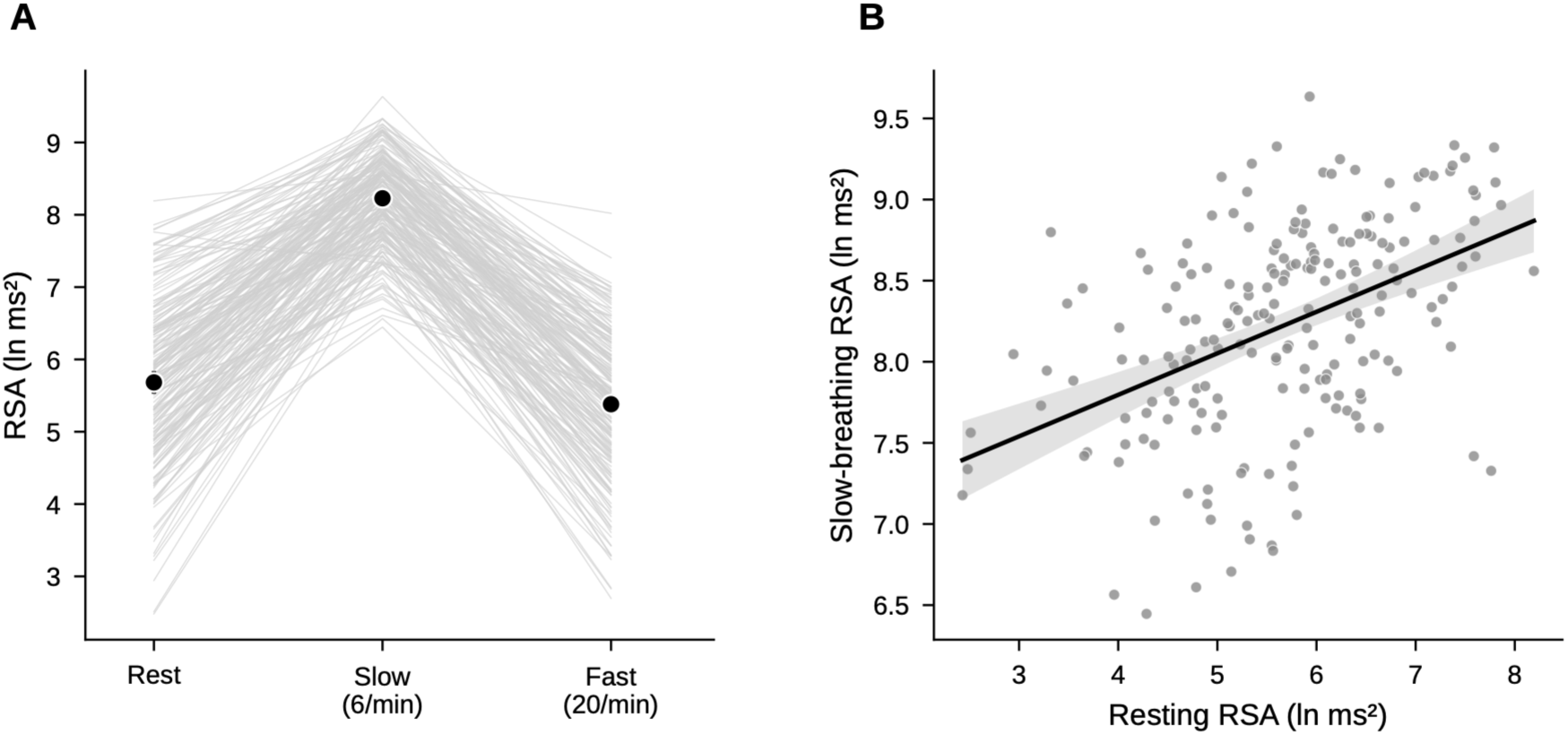
Respiratory Sinus Arrhythmia Across Breathing Conditions. (A) RSA (natural log of power in a band centred on the breathing frequency) at rest, during slow paced breathing (6 breaths/min) and during fast paced breathing (20 breaths/min). Grey lines show individual participants; black points show condition means with 95% confidence intervals (N = 197 with all three conditions). (B) Slow-breathing RSA as a function of resting RSA (r(198) = .46), with the least-squares regression line and its 95% confidence band (N = 200). Rest-adjusted slow-breathing RSA was defined as each participant’s residual from this regression. RSA = respiratory sinus arrhythmia.

Slow-breathing RSA correlated only moderately with resting RSA (r(198) = .46; Fig. 1b), so most between-person variance during slow breathing was not explained by resting RSA. Rest-adjusted slow-breathing RSA was therefore defined as slow-breathing RSA residualized on resting RSA. Heart rate fell slightly during slow breathing (76.5 vs. 78.5 beats/min, p < .001), and heart-rate slowing was only weakly correlated with rest-adjusted slow-breathing RSA (r(198) = .18).

### Generalized anxiety symptoms was associated with lower slow-breathing RSA but not with resting RSA

In a model including resting RSA, age, gender and resting breathing rate (N = 199; Table 2), higher rest-adjusted slow-breathing RSA predicted lower GAD-7 scores (β = −0.21, 95% CI [−0.35, −0.07], p = .004, BF₁₀ = 13.1; Fig. 2a), whereas resting RSA did not (β = −0.09, p = .27, BF₀₁ = 1.8 in this model; without slow-breathing RSA in the model, β = −0.11, p = .18, BF₀₁ = 1.4; Supplementary Table 5). Rest-adjusted slow-breathing RSA explained about 4.6 times as much variance as resting RSA (ΔR² = .041 beyond resting RSA vs. .009 for resting RSA beyond covariates; Fig. 2c). Rest-adjusted fast-breathing RSA was unrelated to GAD-7 scores (β = −0.02, p = .81, BF₀₁ = 2.7). These models show that slow-breathing RSA predicted symptoms beyond resting RSA; whether the association itself differed between conditions was tested separately and is reported below.

**Figure 2.**
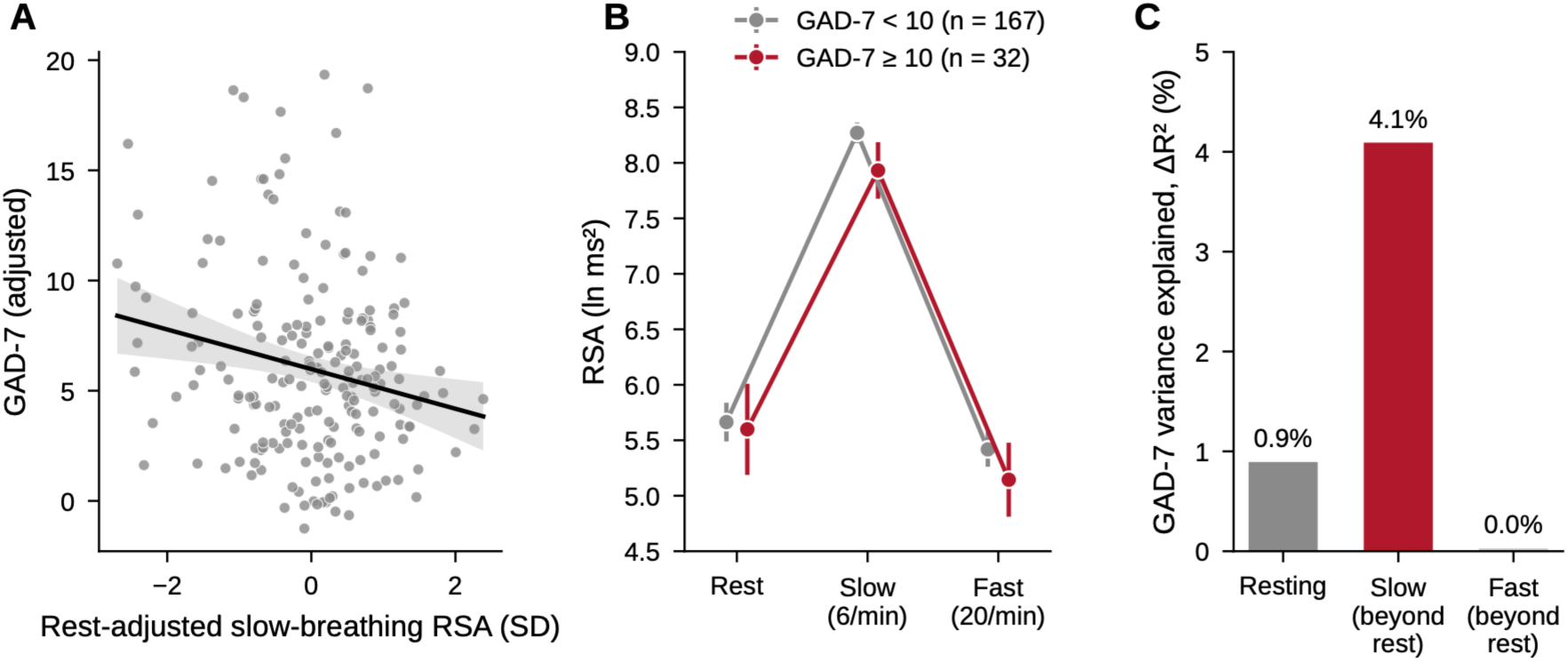
**Generalized Anxiety Is Associated With Lower Slow-Breathing RSA but Not With Resting RSA**. (A) Partial regression plot of GAD-7 scores against rest-adjusted slow-breathing RSA, both adjusted for resting RSA, age, gender and resting breathing rate (β = −0.21, p = .004, N = 199). The line shows the least-squares fit and the shaded band its 95% confidence interval. (B) Mean RSA (95% CI) in each condition for participants below (n = 167) and at or above (n = 32) the threshold for moderate generalized anxiety (GAD-7 ≥ 10). Points are offset horizontally for visibility. (C) Unique variance in GAD-7 scores (ΔR²) explained by resting RSA beyond covariates (from a model without slow-breathing RSA: β = −0.11, p = .18; with rest-adjusted slow-breathing RSA in the model, as in Table 2, β = −0.09, p = .27), by slow-breathing RSA beyond resting RSA (p = .004), and by fast-breathing RSA beyond resting RSA (p = .81). RSA = respiratory sinus arrhythmia; GAD-7 = Generalized Anxiety Disorder-7.

**Table 2.**
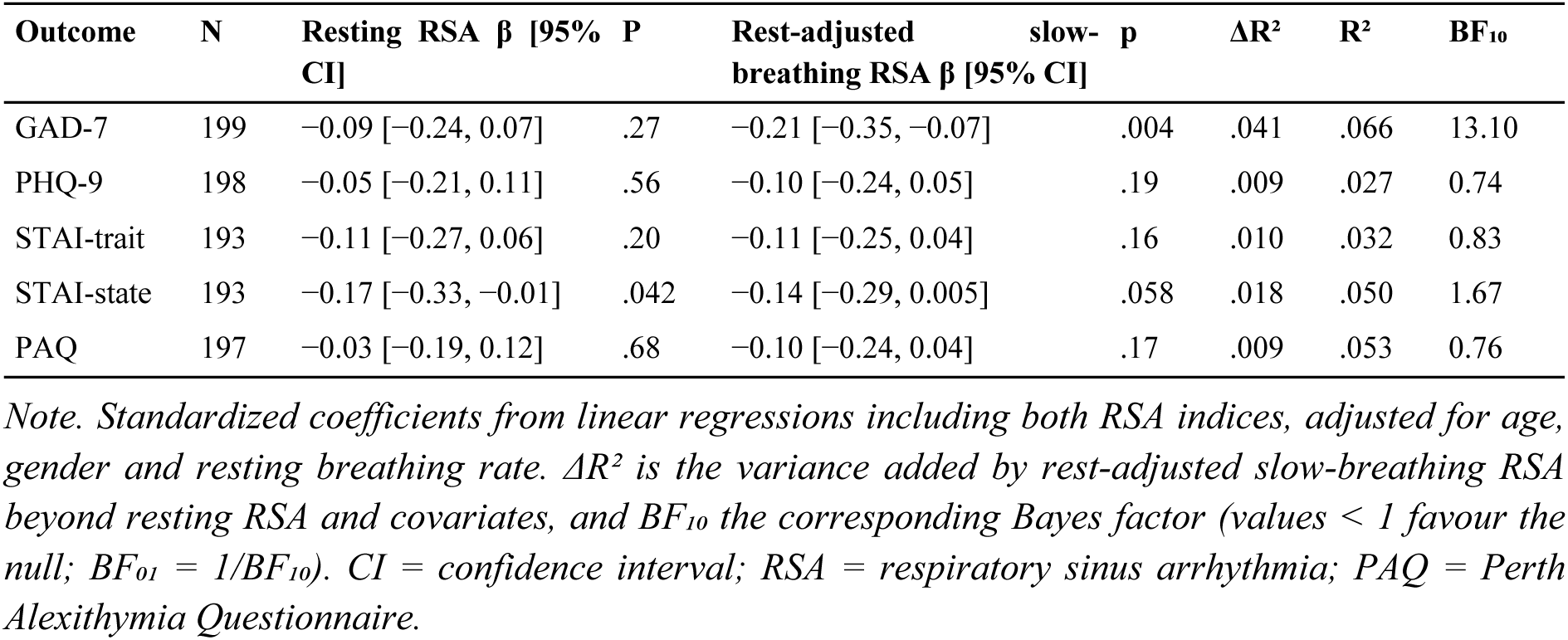
Resting RSA and Rest-Adjusted Slow-Breathing RSA as Simultaneous Predictors of Symptom Outcomes.

Participants with at least moderate generalized anxiety symptoms (GAD-7 ≥ 10; n = 32) had rest-adjusted slow-breathing RSA 0.62 SD lower than those below this threshold (n = 167; 95% CI [−0.99, −0.26], p < .001, BF₁₀ = 43.5). The difference was similar for thresholds from 8 to 12 (−0.40 to −0.69 SD, all ps ≤ .010). The two groups did not differ in resting RSA (5.60 vs. 5.66 ln ms²; −0.04 SD, 95% CI [−0.39, 0.30], p = .80, BF₀₁ = 4.7), but diverged during slow breathing (7.93 vs. 8.27 ln ms²; −0.58 SD, 95% CI [−0.95, −0.21], p = .003, BF₁₀ = 18.4; Fig. 2b). During fast breathing the difference was smaller in SD units and inconclusive (−0.29 SD, p = .15, BF₀₁ = 1.2), although in raw units it was similar to that during slow breathing (−0.27 vs. −0.34 ln ms²), because fast-breathing RSA varied more between participants (SD = 1.01 vs. 0.64; resting RSA, SD = 1.14). Within the mild range (GAD-7 < 10), where scores varied less, the association was weak and non-significant (β = −0.04, p = .65, BF₀₁ = 2.4, n = 167).

In mixed models testing whether the association differed between conditions, the group difference was larger during slow breathing than at rest on the standardized scale (interaction = −0.48, 95% CI [−0.87, −0.08], p = .017) but not in raw units (p = .16), and with continuous GAD-7 scores the corresponding interaction was not significant (standardized: p = .21; raw: p > .99). Relative to fast breathing, the association during slow breathing was stronger for continuous GAD-7 scores on the standardized scale (p = .042) but not for the group comparison (p = .22) or in raw units (ps ≥ .28; Supplementary Table 9). The evidence that the association was specific to slow breathing is therefore moderate.

The difference between the group and continuous analyses reflects the shape of the relation (Supplementary Table 11). Across GAD-7 bands (0–4, 5–9, 10–14 and 15–21), slow-breathing RSA declined steadily (8.33, 8.22, 8.00, 7.77 ln ms²), whereas resting RSA did not (5.87, 5.50, 5.91, 4.92 ln ms²). Participants scoring 10–14 (n = 22) had resting RSA as high as those scoring 0–4, and only the 10 participants scoring 15 or above had low resting RSA. Pooling scores of 10 or above therefore cancelled the resting difference, whereas the continuous resting slope was driven largely by the highest scorers.

### Relationship of RSA to generalized anxiety versus depressive symptoms

Rest-adjusted slow-breathing RSA did not reliably predict depressive symptoms (PHQ-9: β = −0.10, p = .19, BF₀₁ = 1.4), trait or state anxiety (STAI: β = −0.11 and −0.14, ps ≥ .058) or alexithymia (β = −0.10, p = .17; Table 2). Of the five alexithymia subscales, only difficulty identifying negative feelings reached p < .05 (β = −0.15, p = .046, BF₁₀ = 2.0), which would not survive correction for five tests (Supplementary Table 7). Resting RSA was associated only with state anxiety (β = −0.17, p = .042).

Because GAD-7 and PHQ-9 scores were correlated (r(195) = .68), we tested whether the association held with depressive symptoms controlled (n = 197). Rest-adjusted slow-breathing RSA predicted GAD-7 scores with PHQ-9 controlled (β = −0.15, p = .004) but not PHQ-9 scores with GAD-7 controlled (β = 0.06, p = .30). The coefficient for GAD-7 was larger than that for PHQ-9, but this difference did not reach significance (difference = −0.12, 95% bootstrap CI [−0.25, 0.01], p = .064). Participants above the PHQ-9 threshold (n = 31) did not differ from others (−0.25 SD, p = .20, BF₀₁ = 1.5). Because trait anxiety correlated strongly with GAD-7 scores (r(191) = .66) but showed no association, the two were also tested together: rest-adjusted slow-breathing RSA predicted GAD-7 scores with STAI-trait controlled (β = −0.16, p = .005, n = 193) but not STAI-trait scores with GAD-7 controlled (β = 0.04, p = .44). One possible reason for this dissociation is item content: the GAD-7 includes items on difficulty relaxing and restlessness, which the six-item trait scale samples less directly.

### Robustness

The association with GAD-7 scores was robust to adjustment for respiratory amplitude (β = −0.20, p = .007), mean heart period (β = −0.18, p = .014) and both together (β = −0.17, p = .023). It was also robust to alternative outlier treatments; no trimming (β = −0.17, p = .019), winsorizing (β = −0.19, p = .009), rank transformation (β = −0.15, p = .038) and 3.5-MAD exclusion (β = −0.20, p = .005; Supplementary Table 3). No observation was unduly influential (maximum Cook’s distance = 0.07), and excluding the 11 observations with Cook’s distance > 4/n left the effect unchanged (β = −0.18, p = .007)..

In a specification curve crossing six outlier treatments with four covariate sets, the effect was significant in 21 of 24 specifications (β = −0.12 to −0.22; Supplementary Fig. 1). The remaining three, all using rank transformation, were in the same direction but narrowly non-significant (ps ≤ .078), consistent with rank transformation compressing the upper range of GAD-7 scores, where the association was concentrated. No specification predicted PHQ-9 scores (0 of 24). With RMSSD or SDNN in place of RSA, which do not isolate the oscillation at the breathing frequency, the effect was in the same direction but weaker (2 and 3 of 24 specifications significant). Simple difference scores, which do not adjust for resting values, were unrelated to GAD-7 scores (β = −0.06, p = .48), whereas raw slow-breathing RSA was associated with them (β = −0.24, p < .001).

Heart-rate slowing during slow breathing was weakly associated with GAD-7 scores (β = −0.15, 95% CI [−0.29, −0.004], p = .044, BF₁₀ = 2.1), but equally with PHQ-9 scores (β = −0.16, p = .034). With both in the same model, rest-adjusted slow-breathing RSA remained associated with GAD-7 scores (β = −0.18, p = .017) and heart-rate slowing did not (β = −0.12, p = .10; Supplementary Fig. 3).

Condition order (99 participants completed slow breathing first and 100 fast breathing first) did not affect rest-adjusted slow-breathing RSA (difference = −0.18 SD, p = .20) or moderate its association with GAD-7 scores (interaction p = .65; slow first: β = −0.17; fast first: β = −0.24), and did not moderate the association for fast breathing (p = .53). Respiratory amplitude and rate did not differ between participants above and below the GAD-7 threshold in any condition (all ps ≥ .22; slow-breathing amplitude: 6.82 vs. 6.98 a.u.; Supplementary Table 10). The within-person change in respiratory amplitude from rest to slow breathing was also unrelated to GAD-7 scores (r(197) = −.05, p = .45), and adding it to the main model left the association unchanged (β = −0.20, p = .006). At rest, 12 participants breathed below 0.15 Hz; excluding the 11 in the analysis sample, none of whom scored above the GAD-7 threshold, left the association unchanged (β = −0.23, p = .002).

### Cardiorespiratory phase coupling

In further exploratory analyses, we examined phase coupling between heart period and respiration within the respiration-centred band. Higher GAD-7 scores were associated with weaker coupling during slow breathing, adjusted for resting coupling (β = −0.16, 95% CI [−0.29, −0.02], p = .024). This held when PHQ-9 scores (β = −0.24, p = .011) or respiratory amplitude (β = −0.15, p = .026) were controlled. GAD-7 scores were not associated with coupling at rest (β = −0.08, p = .26) or during fast breathing (β = −0.09, p = .17). PHQ-9 scores were unrelated to slow-breathing coupling (β = −0.05, p = .45). Coupling during slow breathing correlated with rest-adjusted slow-breathing RSA (r(197) = .47), and when both were entered together, rest-adjusted slow-breathing RSA remained associated with GAD-7 scores (β = −0.17, p = .047) whereas coupling did not (β = −0.08, p = .36).

### Interoceptive accuracy

Heartbeat counting accuracy average was 0.68 (SD = 0.15, n = 107). In the Heart Rate Discrimination task (n = 91), participants underestimated their heart rate (mean threshold = −6.95 bpm, SD = 14.9; t(90) = −4.46, p < .001), as in previous work (Legrand et al., 2022); the mean absolute threshold was 13.3 bpm (SD = 9.6) and the mean slope 8.5 (SD = 2.6). Neither task was associated with symptoms: GAD-7 (heartbeat counting: β = −0.02, BF₀₁ = 2.7; HRD: β = 0.07, BF₀₁ = 2.2), PHQ-9 (β = −0.12 and 0.00, BF₀₁ = 1.5 and 3.0), or general distress and the specific symptom components (all |β| ≤ 0.13, all ps ≥ .21). None of the 11 self-report scales was associated with either task after correction (all q ≥ .84; Supplementary Table 8). Accuracy was not significantly associated with rest-adjusted slow-breathing RSA (heartbeat counting: β = 0.03, BF₀₁ = 2.5; HRD: β = −0.21, p = .054; see below). Heartbeat counting accuracy was associated with higher resting RSA (β = 0.21, p = .021), but counting accuracy was also higher in participants with lower resting heart rate (r(105) = −.30), and with heart rate controlled the association with resting RSA disappeared (β = 0.06, p = .46). Pooled across tasks, the results were the same (Supplementary Table 9), and rest-adjusted slow-breathing RSA remained associated with GAD-7 scores when pooled accuracy was controlled (β = −0.20, p = .009, BF₁₀ = 7.3, N = 191), with no interaction between the two (p = .24). Two uncorrected associations in the HRD sample, of accuracy with rest-adjusted slow-breathing RSA (p = .054) and of precision with GAD-specific variance (p = .048), did not survive correction across the 16 HRD tests (both q = .38), and the latter was absent for GAD-7 scores themselves (p = .59).

### Self-reported interoception related to general distress, slow-breathing RSA to generalized anxiety beyond general distress

None of the 11 self-reported interoception scores was associated with resting RSA or rest-adjusted slow-breathing RSA after false discovery rate correction (all q ≥ .70; Supplementary Table 2). Several scales were associated with symptoms (Supplementary Table 4), but the association between rest-adjusted slow-breathing RSA and GAD-7 scores remained when each scale was controlled (β = −0.20 to −0.25, all ps ≤ .006) and when all 11 were entered together (β = −0.22, p < .001, N = 196). Self-report and rest-adjusted slow-breathing RSA did not interact (MAIA general score: p = .20; individual scales: all q ≥ .27).

With all 11 scores as one block, self-report uniquely explained 26.8% of the variance in GAD-7 scores and physiology 3.7% (BF₁₀ = 20.5), with 1.1% shared. For PHQ-9 scores, self-report explained a similar share (27.0%), but physiology added almost nothing (0.3%, BF₀₁ = 2.4; Supplementary Fig. 2).

We then decomposed symptom variance into a general distress factor (from GAD-7, PHQ-9 and STAI scores; loadings .70–.94) and GAD- and PHQ-specific residuals, and regressed each on resting RSA, rest-adjusted slow-breathing RSA and self-report (MAIA general score; N = 192; Fig. 3). Self-report was associated with general distress (β = −0.27, BF₁₀ = 180) but not with GAD-specific variance (β = 0.08, BF₀₁ = 1.6). Rest-adjusted slow-breathing RSA showed the reverse pattern; it was associated with GAD-specific variance (β = −0.19, p = .010, BF₁₀ = 6.6) but not reliably with general distress (β = −0.13, BF₁₀ = 1.5). No physiological measure predicted PHQ-specific variance. The same pattern held for most questionnaires (Supplementary Table 6), except that both BPQ scales were also associated with PHQ-specific variance (β = 0.31 and 0.23, BF₁₀ ≥ 29.3), and BPQ body awareness also with GAD-specific variance (β = 0.18, BF₁₀ = 4.4).

**Figure 3.**
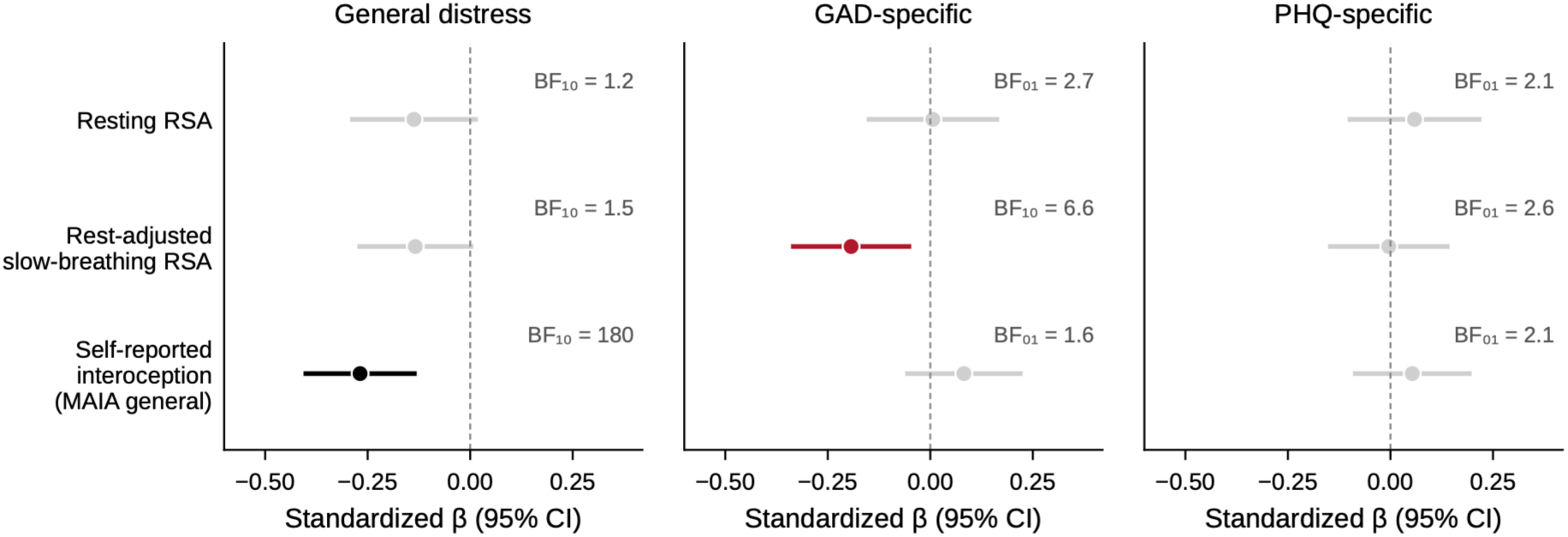
Self-Reported Interoception and Rest-Adjusted Slow-Breathing RSA Relate to Different Components of Symptom Variance. Standardized coefficients with 95% confidence intervals from simultaneous regressions of each symptom component on resting RSA, rest-adjusted slow-breathing RSA and the MAIA general score (mean of six subscales; Ferentzi et al., 2021), adjusted for age, gender and resting breathing rate (N = 192). General distress is the factor score from a one-factor model of GAD-7, PHQ-9, STAI-state and STAI-trait; GAD- and PHQ-specific components are GAD-7 and PHQ-9 scores residualized on this factor. Coloured estimates are significant at p < .05. Results for each questionnaire are given in Supplementary Table 6. BF = Bayes factor; MAIA = Multidimensional Assessment of Interoceptive Awareness.

## Discussion

Generalized anxiety symptoms were related to RSA during slow paced breathing beyond what is predicted by resting RSA, and this effect was largely independent of what participants reported about their own bodies. Participants with at least moderate symptoms had typical resting RSA but lower RSA during slow paced breathing (0.62 SD lower after adjustment for resting RSA). Across the sample, rest-adjusted slow-breathing RSA explained about four to five times as much variance in generalized anxiety symptoms as resting RSA, which showed the weak association typical of this literature. The association held with depressive and trait-anxiety symptoms controlled, although the direct comparison with depressive symptoms was not significant, and it was strongest in participants with at least moderate symptoms. Self-reported interoception explained more symptom variance than RSA did, but this variance was shared across anxiety and depressive symptoms and unrelated to RSA, so the two contributed largely independently to generalized anxiety symptoms. Self-report was not unrelated to physiology altogether, however, it tracked a different component of the cardiac response, heart-rate slowing, which was weakly and equally related to anxiety and depressive symptoms.

The weak association between resting RSA and generalized anxiety symptoms in this sample matches dimensional meta-analytic estimates (r ≈ −.09; Óvári et al., 2026). Case–control comparisons suggest a larger reduction in generalized anxiety disorder (Chalmers et al., 2014). However, that estimate rests on three small studies and may partly reflect antidepressant use in clinical samples (Licht et al., 2009). In addition, like other extreme-group comparisons, they would be expected to exceed dimensional estimates in a non-clinical sample such as ours. Two features of resting measurement may explain why slow breathing was more informative. First, resting RSA depends on spontaneous breathing (Grossman & Taylor, 2007), variance that pacing removes experimentally. Second, resting measures index the vagal state a person happens to be in, whereas slow breathing may assess how far vagal control can be raised under standardized conditions that promote it (Laborde et al., 2018). The findings are compatible with the long-standing proposal that generalized anxiety involves reduced autonomic flexibility (Friedman, 2007; Hoehn-Saric et al., 1989; Thayer & Lane, 2000).

However, it is also possible that slow-breathing RSA may instead provide a cleaner index of tonic vagal function. Several observations are consistent with this account. Firslty, raw slow-breathing RSA correlated with GAD-7 scores at least as strongly as the rest-adjusted index. Secondly, simple difference scores were unrelated to symptoms, and finally, the association was not reliably stronger during slow breathing than in the other conditions. In raw units the anxiety-related difference was similar during slow and fast breathing (−0.34 vs. −0.27 ln ms²), and only some standardized interaction tests were significant. Differences in between-person variance also complicate comparisons across conditions. Slow breathing reduced between-person variation in RSA (SD = 0.64 ln ms², vs. 1.14 at rest and 1.01 during fast breathing), so a given raw difference corresponds to a larger standardized effect during slow breathing. The raw anxiety-related difference was larger during slow breathing than at rest (−0.34 vs. −0.06 ln ms²), but the corresponding raw-unit interactions were not significant, so we cannot fully rule out that this compression of variance contributed to the stronger standardized association during slow breathing. The finding is therefore best described as an association between symptoms and slow-breathing RSA relative to what is predicted by resting RSA, rather than as evidence of a reduced capacity to increase vagal control.

Measurement precision alone, however, does not fully explain the pattern. Spontaneous breathing adds variability to resting RSA, which reduces power, but it should not remove a group difference in mean resting RSA, and participants with at least moderate symptoms had resting RSA close to that of other participants (5.60 vs. 5.66 ln ms²) yet lower RSA during paced breathing. Their spontaneous breathing rate and depth at rest did not differ from those of other participants, so a compensatory breathing pattern at rest is unlikely to have masked a deficit. In exploratory analyses, anxiety was also associated with weaker phase coupling between heart period and respiration during slow breathing, a measure of the timing rather than the size of the oscillation. Two accounts remain. Cardiac vagal outflow in anxious individuals may entrain less tightly to an imposed breathing rhythm. Alternatively, paced breathing, which requires attending to and controlling the breath, may itself be a mild interoceptive challenge to which anxious individuals respond with arousal; heart rate tended to fall less during slow breathing and to rise more during fast breathing in participants above the threshold (ps = .16 and .067). Distinguishing these accounts will require a respiration-controlled baseline outside the resonance range, such as paced breathing at 12 breaths/min, and the time course of RSA and coupling at the onset of paced breathing.

Previous challenge studies in generalized anxiety measured vagal withdrawal under stress, with inconsistent results, and blunted withdrawal has been linked mainly to depression (Adolph et al., 2025; Hamilton & Alloy, 2016; Kircanski et al., 2016). The present findings concern the opposite direction. The two directions may therefore relate differentially to generalized anxiety and depression, which could be tested by combining emotional and respiratory challenges in the same individuals. The clinical deep-breathing test uses the same manoeuvre but did not distinguish individuals with generalized anxiety symptoms from controls when scored against clinical criteria (Kupats et al., 2018). The present results suggest that informative variation lies within the normal range that such criteria do not resolve.

If the association reflects a response to controlled breathing, several mechanisms could contribute. Under the perseverative cognition hypothesis, worry prolongs stress-related physiological activation (Brosschot et al., 2006), so lower slow-breathing RSA may reflect a system habitually held in a state of preparation. Under the contrast avoidance model, individuals with generalized anxiety find relaxation aversive because it exposes them to sharp shifts into negative emotion (Newman & Llera, 2011), and patients with anxiety disorders indeed report more distress than controls after relaxation (Pittig et al., 2013). More proximally, slow breathing may have been uncomfortable for anxious participants, and air hunger or overbreathing would itself alter RSA. Testing these mechanisms will require recording subjective state and end-tidal CO₂ during slow breathing, manipulating worry, and following the measure across treatment.

Interoceptive accuracy was unrelated to symptoms, self-report and rest-adjusted slow-breathing RSA, consistent with Banellis et al. (2026). Its association with resting RSA in the heartbeat counting task was explained by resting heart rate, consistent with the dependence of counting performance on heart rate, and accuracy may become informative only when heartbeats are perceived under challenge. Self-reported interoception, by contrast, explained substantial symptom variance but was unrelated to resting or rest-adjusted slow-breathing RSA, even on the BPQ autonomic reactivity scale. Because this variance was shared across anxiety and depressive symptoms, self-reports may partly index general distress or beliefs about the body rather than bodily function (Desmedt et al., 2022). This parallels the frequent autonomic complaints of individuals with generalized anxiety symptoms, whose autonomic test results did not differ from those of controls (Kupats et al., 2018). Two observations qualify this conclusion. First, the BPQ was additionally associated with variance specific to depressive symptoms, plausibly through overlap with somatic PHQ-9 items. Second, in exploratory analyses, less worry about bodily sensations accompanied greater heart-rate slowing (Supplementary Fig. 3), suggesting that self-report may track coarser bodily changes. Self-report and rest-adjusted slow-breathing RSA nonetheless made largely independent contributions to generalized anxiety symptoms, suggesting that combining them could improve the characterization of anxiety.

Several methodological limitations relating to the methods and the sample need to be acknowledged. Firsty, with regards to the breathing protocol, pacing at a fixed 6 breaths/min rather than at individual resonance frequencies mainly adds noise but could introduce bias if resonance frequency covaries with symptoms or sex. The RSA increase at this rate is amplified by the baroreflex, and sympathetic influences cannot be fully excluded, so the present data cannot separate vagal, baroreflex and sympathetic contributions, although the cardiac arm of the baroreflex is itself predominantly vagal. With regards to the sample, it comprised predominantly female university students with mostly mild symptoms, so replication in clinical populations is needed. Moreover, because participants with self-reported psychiatric disorders were excluded, the sample was non-clinical by design, and the findings cannot be assumed to extend to individuals with diagnosed anxiety disorders.

In conclusion, participants with more severe generalized anxiety symptoms reached lower RSA during slow paced breathing than their resting RSA predicted, although they did not differ at rest. This association was more consistent for generalized anxiety symptoms than for depressive symptoms and was independent of self-reported interoception. Whether it reflects weaker entrainment of vagal outflow to an imposed breathing rhythm or a response to the demands of controlled breathing remains to be determined. Bodily correlates of mental health may nonetheless be more readily identified under standardized physiological conditions than at rest.

## Data Availability

The data for this study are available on the Open Science Framework (https://osf.io/y2tzn/files/qacg3).

## Author contributions

J.S. conceived the study. X.Z. and J.S. developed the methodology. X.Z., H.Q.N. and K.S. collected and curated the data. X.Z, J.S, and T.K.. wrote the analysis code, performed the analyses and prepared the figures. X.Z. and J.S. wrote the original draft, and X.Z., C.T.I., T.K. and J.S. reviewed and edited the manuscript. J.S. supervised and administered the project, provided resources and acquired funding.

## Declaration of Generative AI and AI-assisted technologies in the manuscript preparation process

During the preparation of this work, the authors used Claude to check the grammar of the text. After using this tool/service, the authors reviewed and edited the content as needed and take full responsibility for the content of the published article

## Supplementary Results

**Table S1.** Pearson Correlations Among Key Variables.

| **Variable** | **1** | **2** | **3** | **4** | **5** | **6** | **7** | **8** | **9** | **10** | **11** | **12** | **13** |
|---|---|---|---|---|---|---|---|---|---|---|---|---|---|
| 1. Resting RSA | — |  |  |  |  |  |  |  |  |  |  |  |  |
| 2. Slow-breathing RSA | .46** | — |  |  |  |  |  |  |  |  |  |  |  |
| 3. Rest-adj. slow RSA | −.00 | .89** | — |  |  |  |  |  |  |  |  |  |  |
| 4. GAD-7 | −.12 | −.23** | −.18* | — |  |  |  |  |  |  |  |  |  |
| 5. PHQ-9 | −.05 | −.10 | −.08 | .68** | — |  |  |  |  |  |  |  |  |
| 6. STAI-trait | −.13 | −.15* | −.08 | .67** | .65** | — |  |  |  |  |  |  |  |
| 7. STAI-state | −.16* | −.20** | −.12 | .61** | .52** | .81** | — |  |  |  |  |  |  |
| 8. PAQ | −.04 | −.08 | −.06 | .17* | .16* | .28** | .22** | — |  |  |  |  |  |
| 9. MAIA general | .02 | .03 | .03 | −.14 | −.15* | −.28** | −.23** | −.27** | — |  |  |  |  |
| 10. MAIA Not-Worrying | .08 | .04 | .01 | −.25** | −.23** | −.23** | −.19** | −.20** | .14* | — |  |  |  |
| 11. IAS | .06 | .07 | .04 | −.07 | −.06 | −.21** | −.22** | −.29** | .40** | −.02 | — |  |  |
| 12. BPQ-BA | −.08 | .00 | .08 | .34** | .43** | .26** | .27** | .10 | −.06 | −.28** | −.15* | — |  |
| 13. BPQ-AR | −.17* | −.10 | −.02 | .41** | .45** | .38** | .32** | .28** | −.05 | −.26** | −.18* | .69** | — |
| 14. Age | −.05 | −.11 | −.10 | −.01 | −.13 | −.05 | −.03 | −.12 | .12 | .20** | −.04 | −.16* | −.04 |

*Note.* Pairwise ns = 193–206. Correlations with interoceptive accuracy are given in Supplementary Table 9. RSA = respiratory sinus arrhythmia; PAQ = Perth Alexithymia Questionnaire; MAIA general = mean of six MAIA-2 subscales (Ferentzi et al., 2021); IAS = Interoceptive Accuracy Scale; BPQ-BA = Body Perception Questionnaire body awareness; BPQ-AR = BPQ autonomic reactivity. *p* < *.05.* \* p < .01.

**Table S2.** Interoception Scales Regressed on Resting RSA and Rest-Adjusted Slow-Breathing RSA.

| **Scale** | **Resting RSA β** | **p** | **BF<sub>01</sub>** | **Rest-adj. slow RSA β** | **p** | **BF<sub>01</sub>** |
|---|---|---|---|---|---|---|
| IAS | 0.05 | .55 | 2.4 | 0.04 | .57 | 2.4 |
| BPQ body awareness | −0.11 | .16 | 1.2 | 0.08 | .30 | 1.8 |
| BPQ autonomic reactivity | −0.14 | .074 | 0.71 | −0.03 | .71 | 2.8 |
| MAIA Noticing | −0.11 | .19 | 1.3 | 0.09 | .22 | 1.5 |
| MAIA Not-Distracting | 0.07 | .36 | 2.0 | −0.01 | .93 | 2.8 |
| MAIA Not-Worrying | 0.05 | .53 | 2.6 | 0.04 | .58 | 2.7 |
| MAIA Attention Regulation | −0.02 | .81 | 2.9 | −0.04 | .60 | 2.6 |
| MAIA Emotional Awareness | −0.05 | .57 | 2.5 | 0.08 | .27 | 1.7 |
| MAIA Self-Regulation | 0.06 | .47 | 2.3 | 0.04 | .54 | 2.5 |
| MAIA Body Listening | −0.14 | .088 | 0.79 | 0.06 | .41 | 2.1 |
| MAIA Trusting | 0.11 | .18 | 1.3 | 0.10 | .17 | 1.3 |

*Note.* Standardized coefficients adjusted for age, gender and resting breathing rate (N = 198). Uncorrected p values are shown; after false discovery rate correction across all 22 tests, all qs ≥ .70. BF<sub>01</sub> > 1 favours no association. BPQ autonomic reactivity is the sum of the supradiaphragmatic and subdiaphragmatic subscales.

**Table S3.**
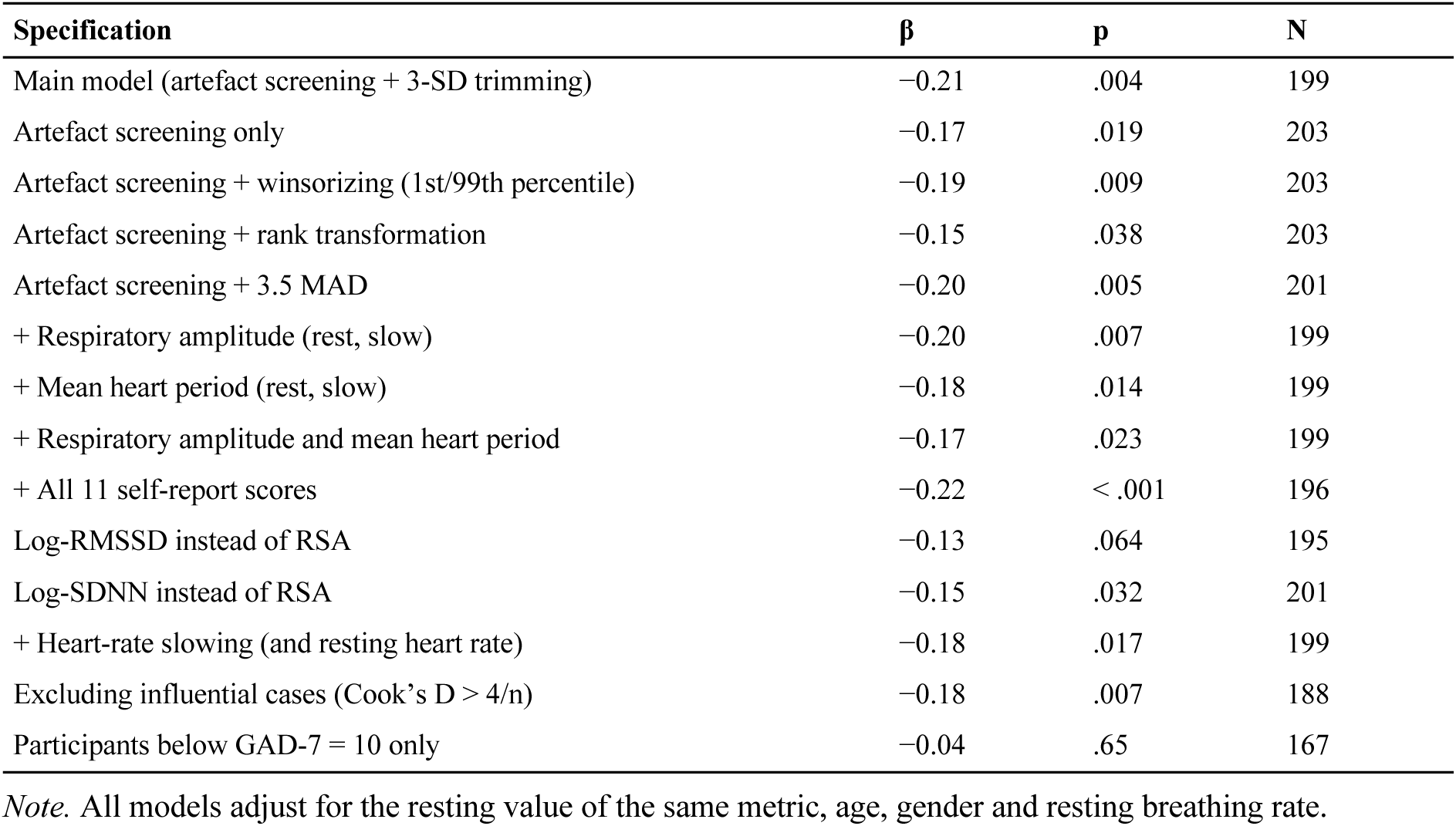
Sensitivity Analyses for the Association Between Rest-Adjusted Slow-Breathing RSA and GAD-7 Scores.

**Table S4.** Associations of Interoception Scales With Symptoms, and the Rest-Adjusted Slow-Breathing RSA Effect Controlling for Each Scale.

| **Scale** | **r with GAD-7** | **r with PHQ-9** | **Scale β on GAD-7** | **p** | **Rest-adj. slow RSA β on GAD-7** | **p** |
|---|---|---|---|---|---|---|
| IAS | −.07 | −.06 | −0.04 | .55 | −0.22 | .003 |
| BPQ body awareness | .34** | .43** | 0.36 | < .001 | −0.25 | < .001 |
| BPQ autonomic reactivity | .41** | .45** | 0.40 | < .001 | −0.21 | .002 |
| MAIA Noticing | −.03 | −.07 | 0.00 | .99 | −0.22 | .003 |
| MAIA Not-Distracting | −.01 | −.06 | −0.01 | .92 | −0.22 | .003 |
| MAIA Not-Worrying | −.25** | −.23** | −0.27 | < .001 | −0.21 | .003 |
| MAIA Attention Regulation | −.16* | −.17* | −0.19 | .008 | −0.23 | .002 |
| MAIA Emotional Awareness | .02 | −.03 | 0.04 | .59 | −0.22 | .003 |
| MAIA Self-Regulation | −.23** | −.18** | −0.23 | .001 | −0.21 | .003 |
| MAIA Body Listening | .01 | .00 | 0.02 | .83 | −0.22 | .003 |
| MAIA Trusting | −.25** | −.24** | −0.23 | .001 | −0.20 | .006 |

*Note.* Correlations are zero-order (ns = 204–205). The final four columns are from separate regressions of GAD-7 scores on each scale together with resting RSA, rest-adjusted slow-breathing RSA, age, gender and resting breathing rate (ns = 197–198).  *p < .05.* \* p < .01.

**Table S5.** Robustness of Key Bayes Factors to the Width of the Prior.

| **Test** | **Index** | **r = 0.354** | **r = 0.5** | **r = 0.707** |
|---|---|---|---|---|
| GAD-7: rest-adjusted slow-breathing RSA beyond resting RSA | BF<sub>10</sub> | 13.1 | 12.7 | 11.2 |
| GAD-7: resting RSA beyond covariates | BF<sub>01</sub> | 1.4 | 1.8 | 2.4 |
| GAD-7: rest-adjusted fast-breathing RSA | BF<sub>01</sub> | 2.7 | 3.6 | 4.8 |
| GAD-7 ≥ 10: lower rest-adjusted slow-breathing RSA | BF<sub>10</sub> | 43.5 | 44.0 | 40.4 |
| GAD-7 ≥ 10: resting RSA | BF<sub>01</sub> | 4.7 | 5.4 | 6.5 |
| PHQ-9: rest-adjusted slow-breathing RSA | BF<sub>01</sub> | 1.4 | 1.7 | 2.2 |
| GAD-specific variance: rest-adjusted slow-breathing RSA | BF<sub>10</sub> | 6.6 | 6.4 | 5.6 |
| General distress: MAIA general score | BF<sub>10</sub> | 180 | 208 | 211 |
| GAD-specific variance: MAIA general score | BF<sub>01</sub> | 1.6 | 1.9 | 2.5 |
| PAQ: rest-adjusted slow-breathing RSA | BF<sub>01</sub> | 1.3 | 1.6 | 2.0 |

*Note.* Default Jeffreys–Zellner–Siow Bayes factors comparing regression models with and without the predictor of interest, adjusted for the same covariates as the corresponding frequentist models, at three prior scales (r). Threshold tests compare participants with GAD-7 scores ≥ 10 with the remainder, adjusting for covariates (and for resting RSA in the test of rest-adjusted slow-breathing RSA). Conclusions were unchanged across prior widths.

**Table S6.** Each Self-Report Measure in Relation to General Distress and Symptom-Specific Variance.

| **Scale** | **General β** | **BF<sub>10</sub>** | **GAD-specific β** | **BF<sub>10</sub>** | **PHQ-specific β** | **BF<sub>10</sub>** | **Rest-adj. slow RSA β (GAD-specific)** |
|---|---|---|---|---|---|---|---|
| MAIA general (6 subscales) | −0.27 | 180 | 0.08 | 0.64 | 0.05 | 0.48 | −0.19 |
| MAIA Not-Worrying | −0.25 | 59.7 | −0.14 | 1.9 | −0.07 | 0.58 | −0.19 |
| MAIA Not-Distracting | −0.09 | 0.68 | 0.12 | 1.1 | 0.03 | 0.41 | −0.19 |
| IAS | −0.18 | 5.6 | 0.14 | 1.9 | 0.11 | 1.1 | −0.20 |
| BPQ body awareness | 0.32 | 1883 | 0.18 | 4.4 | 0.31 | 901 | −0.20 |
| BPQ autonomic reactivity | 0.41 | 1.9 × 10<sup>6</sup> | 0.14 | 1.7 | 0.23 | 29.3 | −0.19 |

*Note.* Each row is a separate set of simultaneous regressions of the symptom component on the scale, resting RSA and rest-adjusted slow-breathing RSA, adjusted for age, gender and resting breathing rate (N = 191–192). General distress is the factor score from GAD-7, PHQ-9, STAI-state and STAI-trait; GAD- and PHQ-specific components are residuals after regression on this factor. The last column gives the coefficient of rest-adjusted slow-breathing RSA for GAD-specific variance in the same model (all ps ≤ .013). BF<sub>10</sub> values < 1 favour the null.

**Table S7.** Resting RSA and Rest-Adjusted Slow-Breathing RSA as Predictors of Alexithymia Subscales.

| **Subscale** | **N** | **Resting RSA β** | **p** | **Rest-adj. slow RSA β** | **p** | **BF<sub>10</sub>** |
|---|---|---|---|---|---|---|
| PAQ N-DIF | 197 | 0.00 | >.99 | −0.15 | .046 | 2.01 |
| PAQ P-DIF | 197 | −0.10 | .23 | −0.06 | .40 | 0.47 |
| PAQ N-DDF | 197 | 0.01 | .92 | −0.09 | .20 | 0.68 |
| PAQ P-DDF | 197 | −0.10 | .19 | −0.01 | .94 | 0.34 |
| PAQ G-EOT | 197 | 0.01 | .86 | −0.09 | .21 | 0.66 |

*Note.* Models as in Table 2. No coefficient survives correction for five subscales.

**Table S8.**
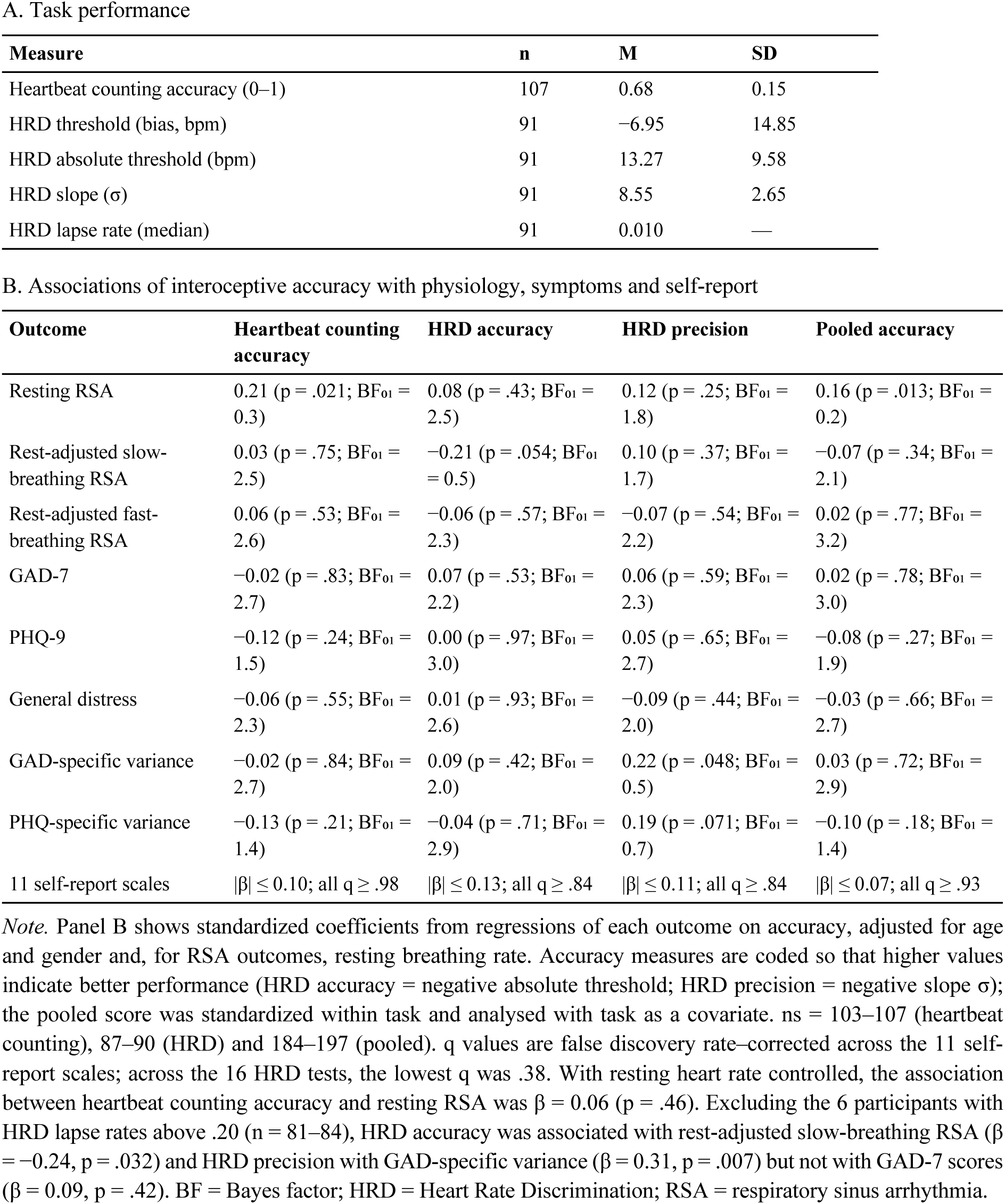
Interoceptive Accuracy: Task Performance and Associations With Physiology, Symptoms and Self-Report.

| A. Task performance |  |  |  |  |
| **Measure** | **n** | **M** | **SD** |  |
|---|---|---|---|---|
| Heartbeat counting accuracy (0–1) | 107 | 0.68 | 0.15 |  |
| HRD threshold (bias, bpm) | 91 | −6.95 | 14.85 |  |
| HRD absolute threshold (bpm) | 91 | 13.27 | 9.58 |  |
| HRD slope (σ) | 91 | 8.55 | 2.65 |  |
| HRD lapse rate (median) | 91 | 0.010 | — |  |
| B. Associations of interoceptive accuracy with physiology, symptoms and self-report |  |  |  |  |
| **Outcome** | **Heartbeat counting accuracy** | **HRD accuracy** | **HRD precision** | **Pooled accuracy** |
| Resting RSA | 0.21 (p = .021; BF<sub>01</sub> = 0.3) | 0.08 (p = .43; BF<sub>01</sub> = 2.5) | 0.12 (p = .25; BF<sub>01</sub> = 1.8) | 0.16 (p = .013; BF<sub>01</sub> = 0.2) |
| Rest-adjusted slow-breathing RSA | 0.03 (p = .75; BF<sub>01</sub> = 2.5) | −0.21 (p = .054; BF<sub>01</sub> = 0.5) | 0.10 (p = .37; BF<sub>01</sub> = 1.7) | −0.07 (p = .34; BF<sub>01</sub> = 2.1) |
| Rest-adjusted fast-breathing RSA | 0.06 (p = .53; BF<sub>01</sub> = 2.6) | −0.06 (p = .57; BF<sub>01</sub> = 2.3) | −0.07 (p = .54; BF<sub>01</sub> = 2.2) | 0.02 (p = .77; BF<sub>01</sub> = 3.2) |
| GAD-7 | −0.02 (p = .83; BF<sub>01</sub> = 2.7) | 0.07 (p = .53; BF<sub>01</sub> = 2.2) | 0.06 (p = .59; BF<sub>01</sub> = 2.3) | 0.02 (p = .78; BF<sub>01</sub> = 3.0) |
| PHQ-9 | −0.12 (p = .24; BF<sub>01</sub> = 1.5) | 0.00 (p = .97; BF<sub>01</sub> = 3.0) | 0.05 (p = .65; BF<sub>01</sub> = 2.7) | −0.08 (p = .27; BF<sub>01</sub> = 1.9) |
| General distress | −0.06 (p = .55; BF<sub>01</sub> = 2.3) | 0.01 (p = .93; BF<sub>01</sub> = 2.6) | −0.09 (p = .44; BF<sub>01</sub> = 2.0) | −0.03 (p = .66; BF<sub>01</sub> = 2.7) |
| GAD-specific variance | −0.02 (p = .84; BF<sub>01</sub> = 2.7) | 0.09 (p = .42; BF<sub>01</sub> = 2.0) | 0.22 (p = .048; BF<sub>01</sub> = 0.5) | 0.03 (p = .72; BF<sub>01</sub> = 2.9) |
| PHQ-specific variance | −0.13 (p = .21; BF<sub>01</sub> = 1.4) | −0.04 (p = .71; BF<sub>01</sub> = 2.9) | 0.19 (p = .071; BF<sub>01</sub> = 0.7) | −0.10 (p = .18; BF<sub>01</sub> = 1.4) |
| 11 self-report scales | \|β\| ≤ 0.10; all q ≥ .98 | \|β\| ≤ 0.13; all q ≥ .84 | \|β\| ≤ 0.11; all q ≥ .84 | \|β\| ≤ 0.07; all q ≥ .93 |

*Note.* Panel B shows standardized coefficients from regressions of each outcome on accuracy, adjusted for age and gender and, for RSA outcomes, resting breathing rate. Accuracy measures are coded so that higher values indicate better performance (HRD accuracy = negative absolute threshold; HRD precision = negative slope σ); the pooled score was standardized within task and analysed with task as a covariate. ns = 103–107 (heartbeat counting), 87–90 (HRD) and 184–197 (pooled). q values are false discovery rate–corrected across the 11 self-report scales; across the 16 HRD tests, the lowest q was .38. With resting heart rate controlled, the association between heartbeat counting accuracy and resting RSA was β = 0.06 (p = .46). Excluding the 6 participants with HRD lapse rates above .20 (n = 81–84), HRD accuracy was associated with rest-adjusted slow-breathing RSA (β = −0.24, p = .032) and HRD precision with GAD-specific variance (β = 0.31, p = .007) but not with GAD-7 scores (β = 0.09, p = .42). BF = Bayes factor; HRD = Heart Rate Discrimination; RSA = respiratory sinus arrhythmia.

**Table S9.** Additional Exploratory Analyses: Condition × Anxiety Interactions, Condition Order, Trait Anxiety and Slow Resting Breathing.

| Analysis | Raw units: estimate [95% CI] | p | Standardized: estimate [95% CI] | p |
|---|---|---|---|---|
| Rest vs. slow × group (GAD-7 ≥ 10) | −0.27 [−0.66, 0.11] | .16 | −0.48 [−0.87, −0.08] | .017 |
| Rest vs. slow × GAD-7 (continuous) | 0.00 [−0.14, 0.14] | >.99 | −0.09 [−0.24, 0.05] | .21 |
| Fast vs. slow × group (GAD-7 ≥ 10) | −0.05 [−0.37, 0.28] | .77 | −0.23 [−0.59, 0.13] | .22 |
| Fast vs. slow × GAD-7 (continuous) | −0.06 [−0.18, 0.05] | .28 | −0.14 [−0.27, −0.01] | .042 |
| Condition order → rest-adjusted slow-breathing RSA |  |  | −0.18 | .20 |
| Order × rest-adjusted slow-breathing RSA (GAD-7) |  |  | 0.06 | .65 |
| Order × GAD-7 (slow-breathing RSA) |  |  | 0.01 | .94 |
| Order × rest-adjusted fast-breathing RSA (GAD-7) |  |  | −0.09 | .53 |
| GAD-7 ~ rest-adj. slow RSA, STAI-trait controlled |  |  | −0.16 | .005 |
| STAI-trait ~ rest-adj. slow RSA, GAD-7 controlled |  |  | 0.04 | .44 |
| Excluding resting breathing < 0.15 Hz |  |  | −0.23 | .002 |

*Note.* Interactions are from linear mixed models with a random intercept per participant, including condition, anxiety, their interaction, age, gender and resting breathing rate (N = 199); standardized models use RSA standardized within condition. A negative interaction indicates a more negative association with anxiety during slow breathing. Order effects are in SD units of the outcome; order was derived from recording timestamps. Other rows are standardized coefficients for rest-adjusted slow-breathing RSA from models as in Table 2.

**Table S10.** Respiration, Heart Rate and Phase Coupling by GAD-7 Group, and Phase Coupling as a Function of GAD-7.

| **Measure** | **GAD-7 < 10** | **GAD-7 ≥ 10** | **p** | **r with GAD-7** |
|---|---|---|---|---|
| Breathing rate, rest (breaths/min) | 14.95 | 15.11 | .81 | .08 |
| Respiratory amplitude, rest (a.u.) | 2.54 | 2.84 | .44 | .02 |
| Breathing rate, slow (breaths/min) | 6.01 | 5.99 | .38 | −.02 |
| Respiratory amplitude, slow (a.u.) | 6.98 | 6.82 | .84 | −.05 |
| Breathing rate, fast (breaths/min) | 19.92 | 19.98 | .22 | .05 |
| Respiratory amplitude, fast (a.u.) | 3.78 | 4.36 | .27 | .06 |
| Heart-rate change, rest to slow (beats/min) | −2.27 | −0.92 | .16 |  |
| Heart-rate change, rest to fast (beats/min) | 1.14 | 3.10 | .067 |  |
| Phase coupling (PLV), rest | 0.83 | 0.80 |  |  |
| Phase coupling (PLV), slow | 0.95 | 0.91 |  |  |
| Phase coupling (PLV), fast | 0.91 | 0.86 |  |  |

*Note.* Group means in the analysis sample (n = 167 and 32); p values from independent-samples t tests. a.u. = arbitrary units (uncalibrated belt). Regression results for phase coupling (Fisher-transformed, adjusted for resting coupling, age, gender and resting breathing rate): slow breathing β = −0.16 (p = .024); with PHQ-9 controlled β = −0.24 (p = .011); fast breathing β = −0.09 (p = .17); rest β = −0.08 (p = .26); PHQ-9 on slow-breathing coupling β = −0.05 (p = .45). PLV = phase-locking value between heart rate and respiration (0 = no synchronization, 1 = perfect synchronization).

**Table S11.** RSA by GAD-7 Band.

| GAD-7 band | n | Resting RSA | Slow-breathing RSA | Fast-breathing RSA | Rest-adjusted slow-breathing RSA |
|---|---|---|---|---|---|
| 0–4 | 76 | 5.87 | 8.33 | 5.40 | 0.06 |
| 5–9 | 91 | 5.50 | 8.22 | 5.44 | 0.04 |
| 10–14 | 22 | 5.91 | 8.00 | 5.22 | −0.28 |
| 15–21 | 10 | 4.92 | 7.77 | 4.97 | −0.26 |

*Note.* Means in the analysis sample. RSA in ln ms²; rest-adjusted slow-breathing RSA is the residual from the regression of slow-breathing on resting RSA (ln ms², mean zero in the full sample).

### Heart-rate slowing and self-reported worry

In exploratory analyses, heart-rate slowing during slow breathing was associated with only one self-report scale after correction for multiple comparisons. Participants whose heart rate slowed more reported less worry about bodily sensations (MAIA Not-Worrying: β = 0.29, 95% CI [0.16, 0.43], q < .001, BF₁₀ = 760). This was robust to respiratory amplitude, RSA and anxiety (β = 0.28, p < .001). Not-Worrying was unrelated to rest-adjusted slow-breathing RSA (β = 0.04, p = .58). Heart-rate slowing was weakly associated with both GAD-7 (β = −0.15, p = .044) and PHQ-9 scores (β = −0.16, p = .034), whereas rest-adjusted slow-breathing RSA was associated only with GAD-7 scores (Supplementary Fig. 3).

**Figure S1.**
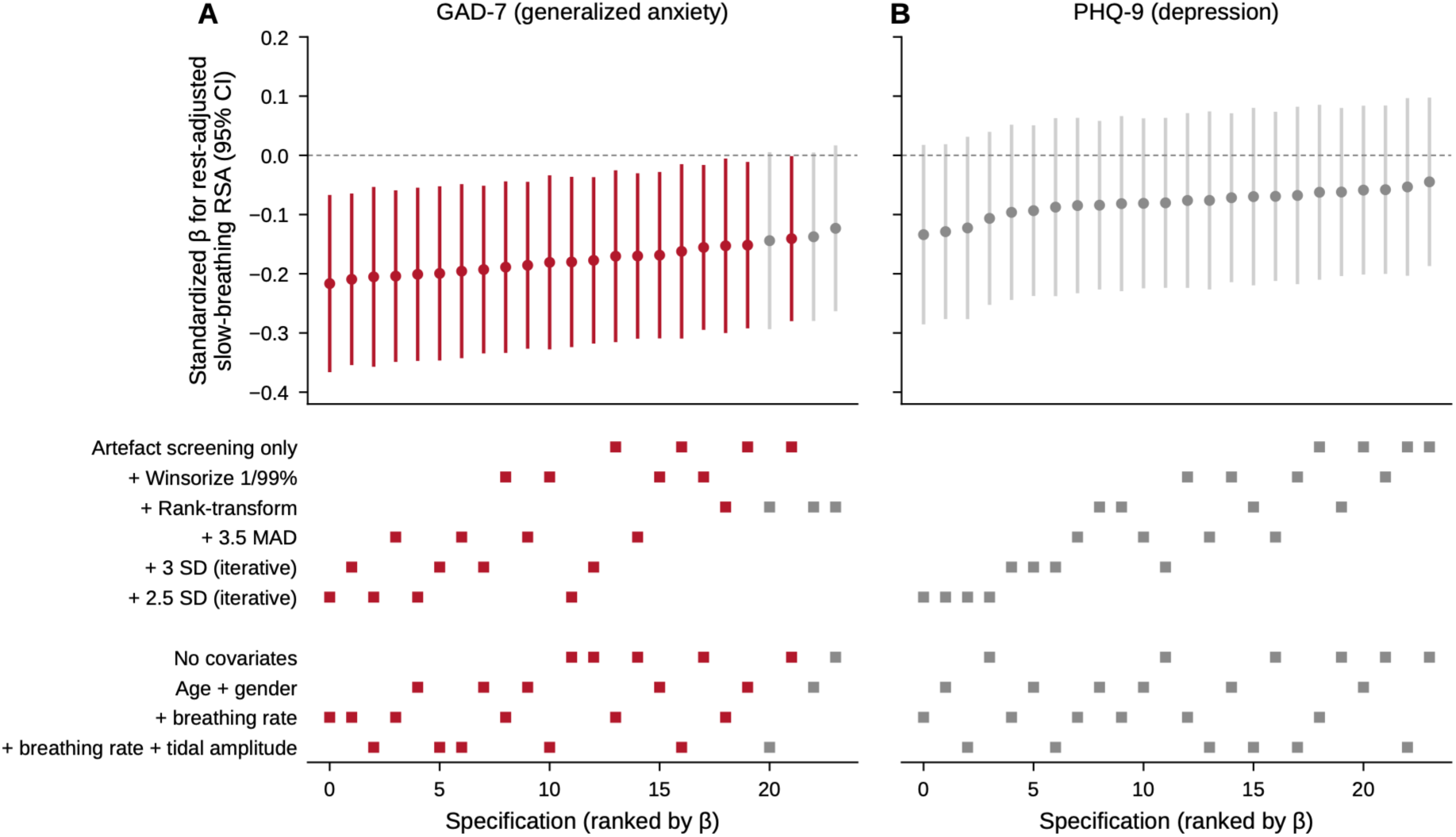
Specification Curve for the Association of Rest-Adjusted Slow-Breathing RSA With Generalized Anxiety and Depression. Each column is one specification (24 per outcome: 6 outlier treatments × 4 covariate sets), ranked by the standardized coefficient for slow-breathing RSA residualized on resting RSA, with resting RSA as a covariate. Upper panels: coefficients with 95% confidence intervals; coloured specifications are significant at p < .05 (GAD-7: 21/24; PHQ-9: 0/24). Lower panels: the analytic choices defining each specification. All specifications apply the artefact screening described in the Methods. Results for RMSSD and SDNN are reported in the text. MAD = median absolute deviation.

**Figure S2.**
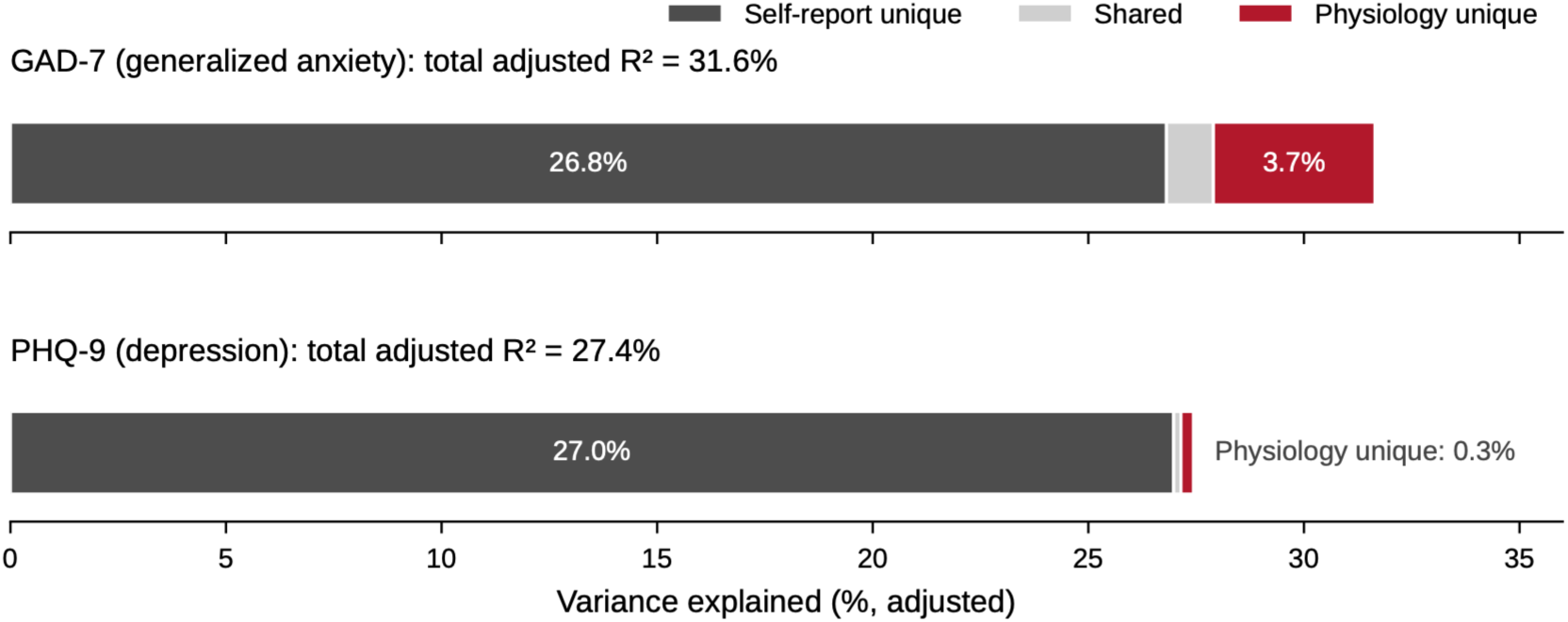
Self-Reported Interoception and Physiology Make Largely Independent Contributions to Symptoms. Partition of the variance in GAD-7 and PHQ-9 scores explained beyond covariates (age, gender, resting breathing rate; N = 196) into variance unique to self-report (all 11 interoception scores), variance unique to physiology (resting RSA and rest-adjusted slow-breathing RSA), and variance shared between the two sets. Values are adjusted R².

**Figure S3.**
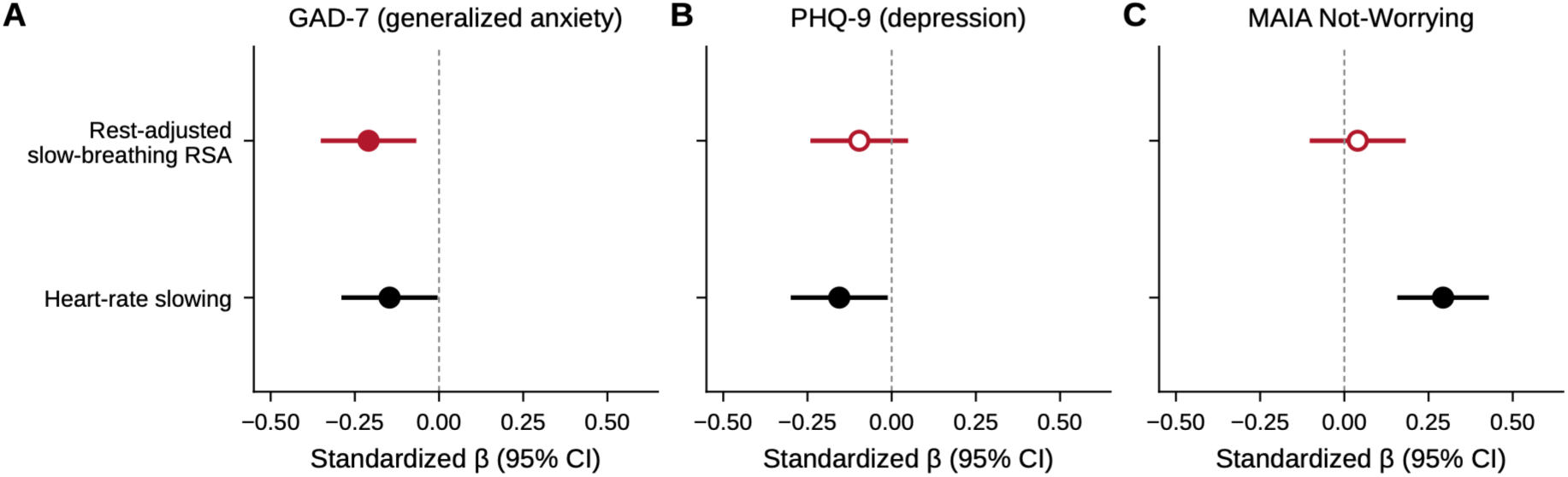
Two Components of the Cardiac Response to Slow Breathing. Standardized regression coefficients with 95% confidence intervals for rest-adjusted slow-breathing RSA (adjusted for resting RSA) and heart-rate slowing (adjusted for resting heart rate; higher values indicate greater slowing), each from a separate model, predicting (A) GAD-7 scores, (B) PHQ-9 scores and (C) MAIA Not-Worrying scores. All models adjust for age, gender and resting breathing rate. Filled markers indicate p < .05. RSA = respiratory sinus arrhythmia; MAIA = Multidimensional Assessment of Interoceptive Awareness.

## References

Adolph, D., Zhang, X. C., Teismann, T., Wannemüller, A., & Margraf, J. (2025). Respiratory sinus arrhythmia—Common and distinct mechanisms of emotional adjustment in the depressive and anxiety disorders spectrum? Psychophysiology, 62(6), Article e70079. 10.1111/psyp.70079

Banellis, L., Nikolova, N., Ehmsen, J. F., Courtin, A. S., Vejlø, M., Tyrer, A., Böhme, R. A., Bavato, F., Hoogervorst, K., Fardo, F., & Allen, M. G. (2026). Interoceptive performance is unrelated to mental health symptoms in a large multi-domain psychophysical investigation. Nature Mental Health, 4(9), 1378–1392. 10.1038/s44220-026-00688-4

Bernstein, D. P., Stein, J. A., Newcomb, M. D., Walker, E., Pogge, D., Ahluvalia, T., Stokes, J., Handelsman, L., Medrano, M., Desmond, D., & Zule, W. (2003). Development and validation of a brief screening version of the Childhood Trauma Questionnaire. Child Abuse & Neglect, 27(2), 169–190. 10.1016/S0145-2134(02)00541-0

Berntson, G. G., Uchino, B. N., & Cacioppo, J. T. (1994). Origins of baseline variance and the law of initial values. Psychophysiology, 31(2), 204–210. 10.1111/j.1469-8986.1994.tb01042.x

Brosschot, J. F., Gerin, W., & Thayer, J. F. (2006). The perseverative cognition hypothesis: A review of worry, prolonged stress-related physiological activation, and health. Journal of Psychosomatic Research, 60(2), 113–124. 10.1016/j.jpsychores.2005.06.074

Cabrera, A., Kolacz, J., Pailhez, G., Bulbena-Cabre, A., Bulbena, A., & Porges, S. W. (2018). Assessing body awareness and autonomic reactivity: Factor structure and psychometric properties of the Body Perception Questionnaire-Short Form (BPQ-SF). International Journal of Methods in Psychiatric Research, 27(2), Article e1596. 10.1002/mpr.1596

Cai, X.-L., Ye, Q., Ni, K., Zhu, L., Zhang, Q., Yin, M., Zhang, Z., Wei, W., Preece, D. A., & Li, B.-M. (2024). Chinese version of the Perth Alexithymia Questionnaire: Psychometric properties and clinical applications. General Psychiatry, 37(2), Article e101281. 10.1136/gpsych-2023-101281

Campbell, A. A., & Wisco, B. E. (2021). Respiratory sinus arrhythmia reactivity in anxiety and posttraumatic stress disorder: A review of literature. Clinical Psychology Review, 87, Article 102034. 10.1016/j.cpr.2021.102034

Chalmers, J. A., Quintana, D. S., Abbott, M. J.-A., & Kemp, A. H. (2014). Anxiety disorders are associated with reduced heart rate variability: A meta-analysis. Frontiers in Psychiatry, 5, Article 80. 10.3389/fpsyt.2014.00080

Desmedt, O., Corneille, O., Luminet, O., Murphy, J., Bird, G., & Maurage, P. (2020). Contribution of time estimation and knowledge to heartbeat counting task performance under original and adapted instructions. Biological Psychology, 154, Article 107904. 10.1016/j.biopsycho.2020.107904

Desmedt, O., Heeren, A., Corneille, O., & Luminet, O. (2022). What do measures of self-report interoception measure? Insights from a systematic review, latent factor analysis, and network approach. Biological Psychology, 169, Article 108289.

Du, Q., Liu, H., Yang, C., Chen, X., & Zhang, X. (2022). The development of a short Chinese version of the State-Trait Anxiety Inventory. Frontiers in Psychiatry, 13, Article 854547.

Ewing, D. J., Martyn, C. N., Young, R. J., & Clarke, B. F. (1985). The value of cardiovascular autonomic function tests: 10 years experience in diabetes. Diabetes Care, 8(5), 491–498. 10.2337/diacare.8.5.491

Ferentzi, E., Olaru, G., Geiger, M., Vig, L., Köteles, F., & Wilhelm, O. (2021). Examining the factor structure and validity of the Multidimensional Assessment of Interoceptive Awareness. Journal of Personality Assessment, 103(5), 675–684. 10.1080/00223891.2020.1813147

Friedman, B. H. (2007). An autonomic flexibility–neurovisceral integration model of anxiety and cardiac vagal tone. Biological Psychology, 74(2), 185–199. 10.1016/j.biopsycho.2005.08.009

Friedman, B. H., & Thayer, J. F. (1998). Anxiety and autonomic flexibility: A cardiovascular approach. Biological Psychology, 47(3), 243–263. 10.1016/S0301-0511(97)00027-6

Garfinkel, S. N., Seth, A. K., Barrett, A. B., Suzuki, K., & Critchley, H. D. (2015). Knowing your own heart: Distinguishing interoceptive accuracy from interoceptive awareness. Biological Psychology, 104, 65–74.

Grossman, P., & Taylor, E. W. (2007). Toward understanding respiratory sinus arrhythmia: Relations to cardiac vagal tone, evolution and biobehavioral functions. Biological Psychology, 74(2), 263–285. 10.1016/j.biopsycho.2005.11.014

Hamilton, J. L., & Alloy, L. B. (2016). Atypical reactivity of heart rate variability to stress and depression across development: Systematic review of the literature and directions for future research. Clinical Psychology Review, 50, 67–79.

Hammel, J. C., Smitherman, T. A., McGlynn, F. D., Mulfinger, A. M. M., Lazarte, A. A., & Gothard, K. D. (2011). Vagal influence during worry and cognitive challenge. *Anxiety, Stress*, & Coping, 24(2), 121–136. 10.1080/10615806.2010.490912

He, X. Y., Li, C., Qian, J., Cui, H. S., & Wu, W. Y. (2010). Reliability and validity of a generalized anxiety disorder scale in general hospital outpatients. Shanghai Archives of Psychiatry, 22(4), 200–203.

Hickman, L., Seyedsalehi, A., Cook, J. L., Bird, G., & Murphy, J. (2020). The relationship between heartbeat counting and heartbeat discrimination: A meta-analysis. Biological Psychology, 156, Article 107949.

Hoehn-Saric, R., McLeod, D. R., & Zimmerli, W. D. (1989). Somatic manifestations in women with generalized anxiety disorder: Psychophysiological responses to psychological stress. Archives of General Psychiatry, 46(12), 1113–1119. 10.1001/archpsyc.1989.01810120055009

Khalsa, S. S., Adolphs, R., Cameron, O. G., Critchley, H. D., Davenport, P. W., Feinstein, J. S., Feusner, J. D., Garfinkel, S. N., Lane, R. D., Mehling, W. E., Meuret, A. E., Nemeroff, C. B., Oppenheimer, S., Petzschner, F. H., Pollatos, O., Rhudy, J. L., Schramm, L. P., Simmons, W. K., Stein, M. B., … Paulus, M. P. (2018). Interoception and mental health: A roadmap. Biological Psychiatry: Cognitive Neuroscience and Neuroimaging, 3(6), 501–513. 10.1016/j.bpsc.2017.12.004

Kircanski, K., Waugh, C. E., Camacho, M. C., & Gotlib, I. H. (2016). Aberrant parasympathetic stress responsivity in pure and co-occurring major depressive disorder and generalized anxiety disorder. Journal of Psychopathology and Behavioral Assessment, 38(1), 5–19. 10.1007/s10862-015-9493-y

Kollai, M., & Kollai, B. (1992). Cardiac vagal tone in generalised anxiety disorder. British Journal of Psychiatry, 161(6), 831–835. 10.1192/bjp.161.6.831

Kroenke, K., Spitzer, R. L., & Williams, J. B. (2001). The PHQ-9: Validity of a brief depression severity measure. Journal of General Internal Medicine, 16(9), 606–613.

Kupats, E., Noviks, I., Vrublevska, J., Kenina, V., Kojalo, U., & Logina, I. (2018). No relationship between generalised anxiety symptoms and cardiovascular autonomic dysfunction. *Neurology*, Psychiatry and Brain Research, 30, 86–90. 10.1016/j.npbr.2018.07.001

Laborde, S., Allen, M. S., Borges, U., Dosseville, F., Hosang, T. J., Iskra, M., Mosley, E., Salvotti, C., Spolverato, L., Zammit, N., & Javelle, F. (2022). Effects of voluntary slow breathing on heart rate and heart rate variability: A systematic review and a meta-analysis. Neuroscience & Biobehavioral Reviews, 138, Article 104711. 10.1016/j.neubiorev.2022.104711

Laborde, S., Mosley, E., & Mertgen, A. (2018). Vagal tank theory: The three Rs of cardiac vagal control functioning—Resting, reactivity, and recovery. Frontiers in Neuroscience, 12, Article 458.

Legrand, N., Nikolova, N., Correa, C., Brændholt, M., Stuckert, A., Kildahl, N., Vejlø, M., Fardo, F., & Allen, M. (2022). The heart rate discrimination task: A psychophysical method to estimate the accuracy and precision of interoceptive beliefs. Biological Psychology, 168, Article 108239. 10.1016/j.biopsycho.2021.108239

Lehrer, P. M., & Gevirtz, R. (2014). Heart rate variability biofeedback: How and why does it work? Frontiers in Psychology, 5, Article 756.

Licht, C. M. M., de Geus, E. J. C., van Dyck, R., & Penninx, B. W. J. H. (2009). Association between anxiety disorders and heart rate variability in The Netherlands Study of Depression and Anxiety (NESDA). Psychosomatic Medicine, 71(5), 508–518. 10.1097/PSY.0b013e3181a292a6

Lin, X.-X., Shen, H.-R., Lin, J.-X., Zhang, Y.-H., Murphy, J., Wang, Y.-Z., Sun, Y.-B., Wang, N., Wang, J.-Y., Wei, G.-X., & Luo, F. (2023). Psychometric validation and refinement of the Chinese Interoceptive Accuracy Scale (IAS) in general population and patients with chronic pain. Journal of Psychosomatic Research, 175, Article 111541. 10.1016/j.jpsychores.2023.111541

Lyonfields, J. D., Borkovec, T. D., & Thayer, J. F. (1995). Vagal tone in generalized anxiety disorder and the effects of aversive imagery and worrisome thinking. Behavior Therapy, 26(3), 457–466.

Mehling, W. E., Acree, M., Stewart, A., Silas, J., & Jones, A. (2018). The Multidimensional Assessment of Interoceptive Awareness, Version 2 (MAIA-2). PLOS ONE, 13(12), Article e0208034.

Murphy, J., Brewer, R., Plans, D., Khalsa, S. S., Catmur, C., & Bird, G. (2020). Testing the independence of self-reported interoceptive accuracy and attention. Quarterly Journal of Experimental Psychology, 73(1), 115–133.

Newman, M. G., & Llera, S. J. (2011). A novel theory of experiential avoidance in generalized anxiety disorder: A review and synthesis of research supporting a contrast avoidance model of worry. Clinical Psychology Review, 31(3), 371–382. 10.1016/j.cpr.2011.01.008

Nunan, D., Sandercock, G. R. H., & Brodie, D. A. (2010). A quantitative systematic review of normal values for short-term heart rate variability in healthy adults. Pacing and Clinical Electrophysiology, 33(11), 1407–1417. 10.1111/j.1540-8159.2010.02841.x

Oliveira, M., da-Silva, M., Carvalho, L., Ribeiro-Carreira, A., Pereira, A. R., Sampaio, A., Coutinho, J., & González-Villar, A. J. (2025). Interplay between cortical and neurocardiac interoceptive processes and its association with self-reported interoceptive sensibility. Brain Topography, 38(4), Article 48. 10.1007/s10548-025-01122-1

Óvári, T., Zabihi, A. Á., Galdzytska, N., Engh, M. A., Veres, D. S., Paczkó, M., Ferdinandy, P., Papp, R. E., Hegyi, P., Alotti, N., & Duray, G. (2026). Heart rate variability and anxiety severity: A systematic review and meta-analysis. Comprehensive Psychiatry, 150, Article 152738. 10.1016/j.comppsych.2026.152738

Pittig, A., Arch, J. J., Lam, C. W. R., & Craske, M. G. (2013). Heart rate and heart rate variability in panic, social anxiety, obsessive–compulsive, and generalized anxiety disorders at baseline and in response to relaxation and hyperventilation. International Journal of Psychophysiology, 87(1), 19–27. 10.1016/j.ijpsycho.2012.10.012

Porges, S. W. (1993). Body Perception Questionnaire. Laboratory of Developmental Assessment, University of Maryland.

Preece, D., Becerra, R., Robinson, K., Dandy, J., & Allan, A. (2018). The psychometric assessment of alexithymia: Development and validation of the Perth Alexithymia Questionnaire. Personality and Individual Differences, 132, 32–44. 10.1016/j.paid.2018.05.011

Quigley, K. S., Gianaros, P. J., Norman, G. J., Jennings, J. R., Berntson, G. G., & de Geus, E. J. C. (2024). Publication guidelines for human heart rate and heart rate variability studies in psychophysiology—Part 1: Physiological underpinnings and foundations of measurement. Psychophysiology, 61(9), Article e14604. 10.1111/psyp.14604

Rouder, J. N., & Morey, R. D. (2012). Default Bayes factors for model selection in regression. Multivariate Behavioral Research, 47(6), 877–903.

Schandry, R. (1981). Heart beat perception and emotional experience. Psychophysiology, 18(4), 483–488. 10.1111/j.1469-8986.1981.tb02486.x

Shields, R. W., Jr. (2009). Heart rate variability with deep breathing as a clinical test of cardiovagal function. Cleveland Clinic Journal of Medicine, 76(4, Suppl. 2), S37–S40. 10.3949/ccjm.76.s2.08

Spielberger, C. D., Gorsuch, R. L., Lushene, R., Vagg, P. R., & Jacobs, G. A. (1983). Manual for the State-Trait Anxiety Inventory. Consulting Psychologists Press.

Spitzer, R. L., Kroenke, K., Williams, J. B., & Löwe, B. (2006). A brief measure for assessing generalized anxiety disorder: The GAD-7. Archives of Internal Medicine, 166(10), 1092–1097.

Teng, B., Wang, D., Su, C., Zhou, H., Wang, T., Mehling, W. E., & Hu, Y. (2022). The Multidimensional Assessment of Interoceptive Awareness, Version 2: Translation and psychometric properties of the Chinese version. Frontiers in Psychiatry, 13, Article 970982.

Thayer, J. F., Friedman, B. H., & Borkovec, T. D. (1996). Autonomic characteristics of generalized anxiety disorder and worry. Biological Psychiatry, 39(4), 255–266.

Thayer, J. F., & Lane, R. D. (2000). A model of neurovisceral integration in emotion regulation and dysregulation. Journal of Affective Disorders, 61(3), 201–216.

Wang, N., Ren, F., & Zhou, X. (2020). Factor structure and psychometric properties of the Body Perception Questionnaire–Short Form (BPQ-SF) among Chinese college students. Frontiers in Psychology, 11, Article 1355. 10.3389/fpsyg.2020.01355

Wang, W., Bian, Q., Zhao, Y., Li, X., Wang, W., Du, J., Zhang, G., Zhou, Q., & Zhao, M. (2014). Reliability and validity of the Chinese version of the Patient Health Questionnaire (PHQ-9) in the general population. General Hospital Psychiatry, 36(5), 539–544.

Wilder, J. (1967). Stimulus and response: The law of initial value. John Wright & Sons.

Zamariola, G., Maurage, P., Luminet, O., & Corneille, O. (2018). Interoceptive accuracy scores from the heartbeat counting task are problematic: Evidence from simple bivariate correlations. Biological Psychology, 137, 12–17.

Zhao, X. F., Zhang, Y. L., Li, L. F., & Zhou, Y. F. (2005). Evaluation on reliability and validity of Chinese version of childhood trauma questionnaire. Chinese Journal of Clinical Rehabilitation, 9(16), 209–211.

